# The mutational landscape defines the proteome and spatial organization of tumor, stroma and immune cells in ovarian cancer

**DOI:** 10.1101/2025.06.01.655822

**Authors:** Kruttika Dabke, Nicole Gull, Simion Kreimer, Pei-Chen Peng, Michael A. Diaz, Bassem Ben Cheikh, Amisha Dhawan, Katrina Evans, Nathan Ng, Xioapu Yuan, Bobbie J. Rimel, Andrew J. Li, Beth Y. Karlan, Simon V. Knott, Simon A. Gayther, Sarah Parker, Michelle R. Jones

## Abstract

High-grade serous ovarian cancer (HGSOC) is a highly aggressive and lethal form of ovarian cancer. Challenges to diagnosis and treatment include a lack of effective screening methods for early detection, the absence of cancer-specific symptoms, and the development of chemoresistance. The genomic instability of HGSOC, further complicated by homologous recombination deficiency (HRD), leads to heterogeneity in HGSOC tumors and patient response to treatment. This makes it challenging to develop a single, effective diagnostic and treatment approach for this disease. Proteogenomic studies have provided some insight into HGSOC biology, but a deeper understanding of the tumor proteome through chemotherapy and disease recurrence is needed. Here we have profiled the proteome of tumors from 29 HGSOC patients before and after multiple rounds of chemotherapy. The proteome of HGSOC tumors remained unchanged for individual patients even after numerous rounds of chemotherapy. Differential expression analysis revealed known and novel proteins associated with chemoresistance and HRD status, further supported by similar changes at the genetic and epigenetic levels. We found that HRD affected proteins related to immune pathways. HRD was also associated with more shared T cell receptor (TCR) CDR3 repertoires of tumor-infiltrating T cells and increased neoantigen counts. ERAP1, a protein involved in peptide trimming before antigen presentation to immune cells via MHC-class I, was overexpressed in homologous recombination proficient (HRP) tumors and negatively correlated to neoantigen count. Its potential role in immune suppression in HRP tumors makes it an attractive therapeutic target that may be effective in combination with immunotherapy, particularly for HRP tumors. 26-plex immunostaining of HGSOC whole tissues further revealed significant differences in the spatial proximities of immune cells to each other and to tumor cells based on HRD status. Through proteomic and imaging analysis, this work has shown that the immune landscape of HGSOC tumors is influenced by homologous recombination status and identified candidate drivers of HGSOC biology.

## Introduction

Ovarian cancer is the second most common cause of gynecological cancer death, where 1 in 6 women die within the first 90 days of diagnosis. The lack of reliable markers for early detection leading to diagnostic delay and resistance to chemotherapy accounts for this disease’s high morbidity and mortality ^1^. High-grade serous ovarian cancer (HGSOC) is the most common histological subtype, accounting for more than 70% of cases of epithelial ovarian cancer ^2^. Most HGSOC cases are sporadic, but 15-20% of women have inherited highly penetrant deleterious variants in *BRCA1* and *BRCA2* genes or moderately penetrant variants in other genes from the homologous recombination repair pathway ^3,4^. Driver mutations in *TP53* are ubiquitous in HGSOC, which in combination with *BRCA1/2* loss lead to chromosomal instability and the widespread accumulation of copy number alterations ^5^. Other genomic changes, like *RB1*, *NF1*, *PTEN* loss or *CCNE1* amplification, and somatic *BRCA1/2* mutations, occur in various combinations and lead to extensive genomic heterogeneity among HGSOC tumors ^6,7^. Homologous recombination deficiency (HRD) is a crucial determinant of platinum sensitivity in HGSOCs, patients with HRD,through either germline variants or somatic loss, have increased survival and response to platinum and taxane based therapies ^8,9^. In recent years the use of inhibitors of poly (ADP ribose) polymerase (PARP), which is recruited to the site of single strand breaks in DNA to mediate repair, in HRD tumors has improved survival in such HGSOC cases ^10^.

Previous studies have established that significant differences in tumor immune microenvironment profiles may contribute to differences in survival in patients with HRD tumors ^11^. The cyclic GMP-AMP synthase-stimulator of interferon genes-signal transducer and activator of transcription pathway (cGAS-STING-STAT) that is activated in response to the sensing of double stranded DNA as an antiviral immune response was recently reported as upregulated in HRD tumors ^12^ and data from animal models and ovarian cancer cell lines support this ^13^. *BRCA1* mutated tumors show an increase in cGAS-STING mediated intratumoral T-cell infiltration ^14^, increased CD8+ tumor infiltrating lymphocytes (TILs) ^15^, and increased VEGFA levels ^16^. STING expression has also been shown to mediate an anti-tumor immune response in the presence of PARP inhibition therapy underscoring the importance of this pathway in treatment response ^17,18^. Despite advances in understanding the mechanisms behind immune response in HGSOC tumors the outcomes of immunotherapy trials have been underwhelming. Several clinical trials testing checkpoint inhibitors in ovarian cancer have been discontinued due to negative outcomes ^19–21^, but combination studies of immune therapies with other therapies are underway. Immunotherapy continues to be challenging in HGSOC, with additional studies needed to identify potentially sensitive populations or a means of sensitizing individuals to immunotherapy.

The genome, transcriptome, and proteome of chemotherapy naïve or neoadjuvant treated HGSOC tumors have been well studied. Large consortia have assayed the genomic and transcriptomic alterations associated with disease and their functional consequences ^4,22,23^. Proteogenomic characterization of chemotherapy naïve HGSOC tumors has uncovered drivers of HGSOC biology ^24,25^. Proteins and pathways associated with changes in the tumor genome and proteome during the development of chemoresistance have been inferred from patient-derived cell lines or ascites samples in previous studies ^26–29^. Candidate prognostic biomarkers have been identified using large scale genetic, proteomic and plasma proteomic data ^30–32^. However, the proteome of HGSOC patient-derived tumors has yet to be extensively studied in the context of chemoresistance and disease relapse, and homologous recombination deficiency. Previous studies of the HGSOC proteome have focused primarily on chemotherapy naïve or neoadjuvant tumors with resistance inferred from disease-free survival or focused on ascites or proximal liquid biopsies, which do not truly reflect the tumor proteome after chemotherapy. An improved understanding of the dysregulated proteome and subsequent immune response in HGSOC recurrence can facilitate enhanced mechanistic understanding for adjuvant drug design and the development of clinical assays, which can be rapidly implemented to improve patient diagnosis and treatment. To this end, we performed a proteogenomic analysis of paired chemotherapy naïve (primary) and chemoresistant (recurrent) HRD and HRP tumors from women with extensive clinical annotations available to map the proteomic markers and spatially resolve the tumor-immune landscape through disease recurrence in HGSOC.

## Results

### The proteomic landscape of high-grade serous ovarian cancer

We assessed the global protein expression profiles of paired primary and recurrent tumor samples and tumor-adjacent stromal samples from 29 patients (**Supp Table 1**). Fifteen patients were classified as HRD and seventeen as HRP based on germline *BRCA1/2* deleterious variants and somatic promoter hypermethylation of *BRCA1* and *RAD51C* (**Supp Table 1 & 2**). We used data independent acquisition mass spectrometry (DIA-MS) to profile the proteome of samplings of the epithelial enriched tumor bed and stroma compartments from women with HGSOC (**Figure 1A; Supp Table 2**). These tumors were previously profiled for genome-wide methylation using whole genome bisulfite sequencing (WGBS) and gene expression using RNA-seq ^33^.

**Figure 1:**
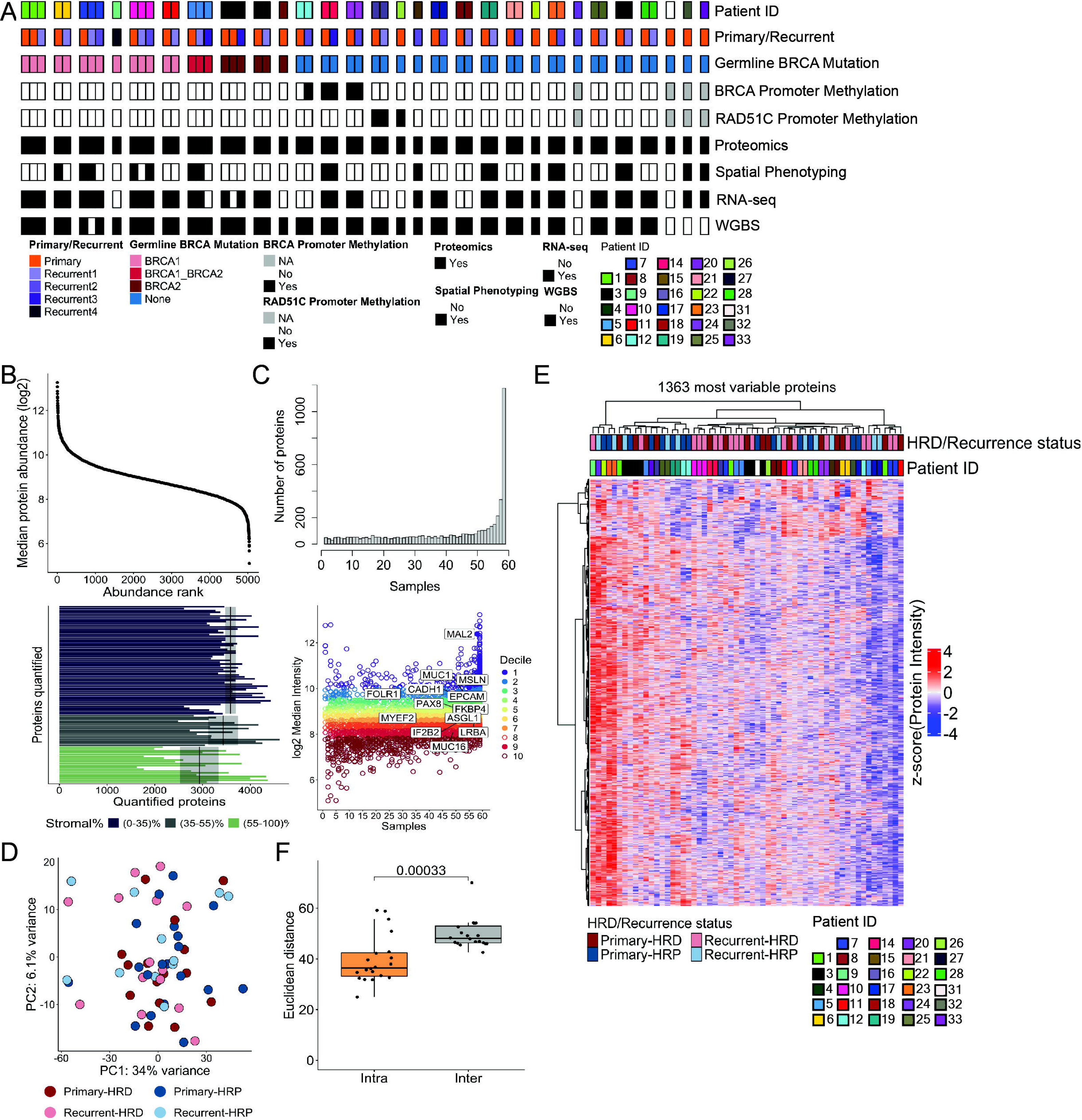
The tumor proteome is stably conserved throughout disease relapse. A. Schematic of tumor samples profiled for proteomics, Phenocycler spatial phenotyping, whole genome bisulfite sequencing, and RNA-seq with clinical and genetic information. B. (Top) Proteins ranked by median abundance levels across 96 samples. (Bottom) The total number of proteins quantified in each sample; (0-35%-Tumor, 35-55%-Mix, 55-100%-Stroma). Black vertical lines indicate the mean per group with a 95% confidence interval shown as a gray-shaded area around the mean. C. (Top) The bar plot on top shows the total number of proteins quantified in 59 tumor samples. (Bottom) Distribution of median log2 protein intensities across the 59 tumor samples against the number of samples they were detected in, colored by decile (1= most abundant proteins and 10 = least abundant proteins). D. Protein expression driven principal component analysis reveals sample heterogeneity in the context of clinical and genomic features. E. Hierarchical clustering of tumors using the top 50% of most variable proteins (n = 1365) is mainly driven by patient IDs instead of recurrence or homologous recombination deficiency status. F. Intra-patient Euclidean distance measured after hierarchical clustering (E) is significantly shorter for pairs of primary and recurrent tumors indicating similar proteomic profiles between primary and recurrent tumors from the same patient (intra-patient vs inter-patient).

We identified 5031 proteins within our sample group, which was made up of 59 tumor samples from 29 patients (**Supp Table 3**). The median dynamic range of protein abundance in the log2 scale spanned 5.09-13.26 (**Figure 1B; top panel**). Protein expression levels across individual tumors in the cohort were highly correlated (mean Pearson’s r value between all samples was 0.66), with a within-sample mean of 0.95 observed across technical replicates (**Supp Figure 1A**). Correlation of expression was similar when measured within only tumor and stroma samples (0.68 and 0.67, respectively) across the cohort. Pooled samples run across multiple mass spectrometry batches clustered together, indicating minimal batch effects (**Supp Figure 1A**). To avoid adverse effects of cellular admixtures on downstream data analysis, relative neoplastic and stromal content within each sample was assigned based on classification generated using machine learning tools applied to H&E stained images and protein expression profiles, assigning samples to one of three groups; tumor, stroma or mix, ^34^ (**Supp Figure 1B,C**). An average of 3585 proteins were quantified within each tumor sample, 3430 proteins within each mixed sample, and 2932 proteins within each stromal sample (**Figure 1B, bottom panel; Supp Table 4**). There was improved clustering of tumor and stromal samples after re-classification based on unsupervised hierarchical clustering using proteins (obtained from publicly available data ^24^) that characterized tumor and stromal components from the same tissue (**Supp Figure 1D, Supp Table 5**).

Differential expression analysis between tumor and stromal samples (those assigned a mixed phenotype were excluded from this analysis) revealed 864 proteins upregulated and 269 proteins downregulated in tumor samples (log2FC ± 0.5, FDR < 0.01, **Supp Figure 1E; Supp Table 6**). Differentially expressed proteins between tumor and stroma detected in our cohort largely replicated previously reported results generated with an alternative proteomics methodology on a separate cohort of patients ^24^ (**Supp Table 6**, overlap of 84% upregulated in tumor and 72% of proteins upregulated in stromal samples as determined by log2FC ± 0.5, FDR < 0.01). Proteins upregulated in tumor samples were enriched in mitochondrial biogenesis, DNA double strand break repair, rRNA processing, and translation pathways. Proteins upregulated in the stromal samples were enriched in complement cascade, extracellular matrix organization (ECM), and integrin cell surface interaction pathways (FDR < 0.05; **Supp Figure 1F, Supp Table 7**).

Of the 5031 proteins we quantified across all 59 tumor samples, 1176 were quantified in every sample (**Figure 1C, top panel; Supp Table 3**). Annexin A2, the most abundant protein identified in all samples in our cohort, is known to be overexpressed in ovarian cancer and is an independent prognostic marker of survival for ovarian cancer patients ^35^ (**Supp Table 3**). We grouped the proteins into deciles based on their median abundance, with the most abundant proteins in the first decile (**Figure 1C, bottom panel**). More abundant proteins were detected across a larger proportion of the samples than those expressed in lower deciles, as has been observed in previous studies ^36^. Proteins in the top decile were enriched in pathways associated with housekeeping functions like mRNA processing, ribonucleoprotein complex biogenesis, generation of precursor metabolites, and energy. Of the top 10 proteins in decile 1, citrate synthase and prohibitin 1 are known to be involved in stemness and chemoresistance in ovarian cancer ^37–39^. Proteins in the bottom decile were enriched in pathways like dephosphorylation, regulation of small GTPase, and positive regulation of DNA metabolic processes (GO term-Biological processes; **Supp Figure 2A; Supp Table 8**). Proteins associated with ribosomal biogenesis were stably expressed across tumor samples (mean SD = 0.55) while previously reported proteins overexpressed in ovarian cancer were observed across the spectrum of expression intensity, with a range of variability across tumor samples (SD range – 0.56 to 1.75; **Supp Figure 2B; Supp Table 9**). This was consistent with other mass spectrometry data from ovarian cancer cell lines ^40^.

Protein - transcript correlations were performed across 41 common samples between the RNA-seq and proteomic data sets. Protein expression correlated well to gene expression (measured by total RNA-Seq) with a median Spearman correlation of 0.34, where 1403 gene-protein pairs (44%) were positively correlated at FDR < 0.05, which is similar to findings from other proteogenomic studies ^22^ ^41^ ^42^ ^43^ (**Supp Figure 2C**). A stronger correlation was observed for more dynamic proteins known to be transcriptionally regulated in lieu of nutrient demand or other distress, e.g., Interferon gamma response, CD40 signaling, EGFR signaling, and keratinization. In comparison, a weaker correlation was observed for more stable and abundant proteins with housekeeping functions: e.g., mRNA splicing, complex 1 biogenesis, and small nuclear ribonucleoproteins (snRNPs) assembly (**Supp Table 10**). The expected distribution of protein abundance across samples, lack of batch effects, and improved clustering of tumor and stromal samples after re-classification with the stromal score were indicative of the high quality of the proteomics data set.

### The tumor proteome is stably conserved throughout disease relapse

Principal component analysis performed with proteins quantified in more than 80% of the samples (n=2729 proteins) highlighted sample heterogeneity in the context of recurrence and HRD status (**Figure 1D**). Principal components analysis showed that some patients’ primary tumors clustered closely with their recurrent tumors, while others were farther apart along the first and second principal components (**Supp Figure 2D,E**). There were no clear clusters that indicated widely shared protein expression programs based on primary/recurrent or HRD status. We performed semi-supervised hierarchical clustering using the top 50% most variable proteins (n=1365) quantified across the 59 tumor samples. Primary tumors were significantly closer in Euclidean distance to recurrent tumors from their respective donor than tumors from other donors (shorter intra-patient distance than inter-patient distance) (**Figure 1E,F; Supp Figure 2F**).

**Figure 2:**
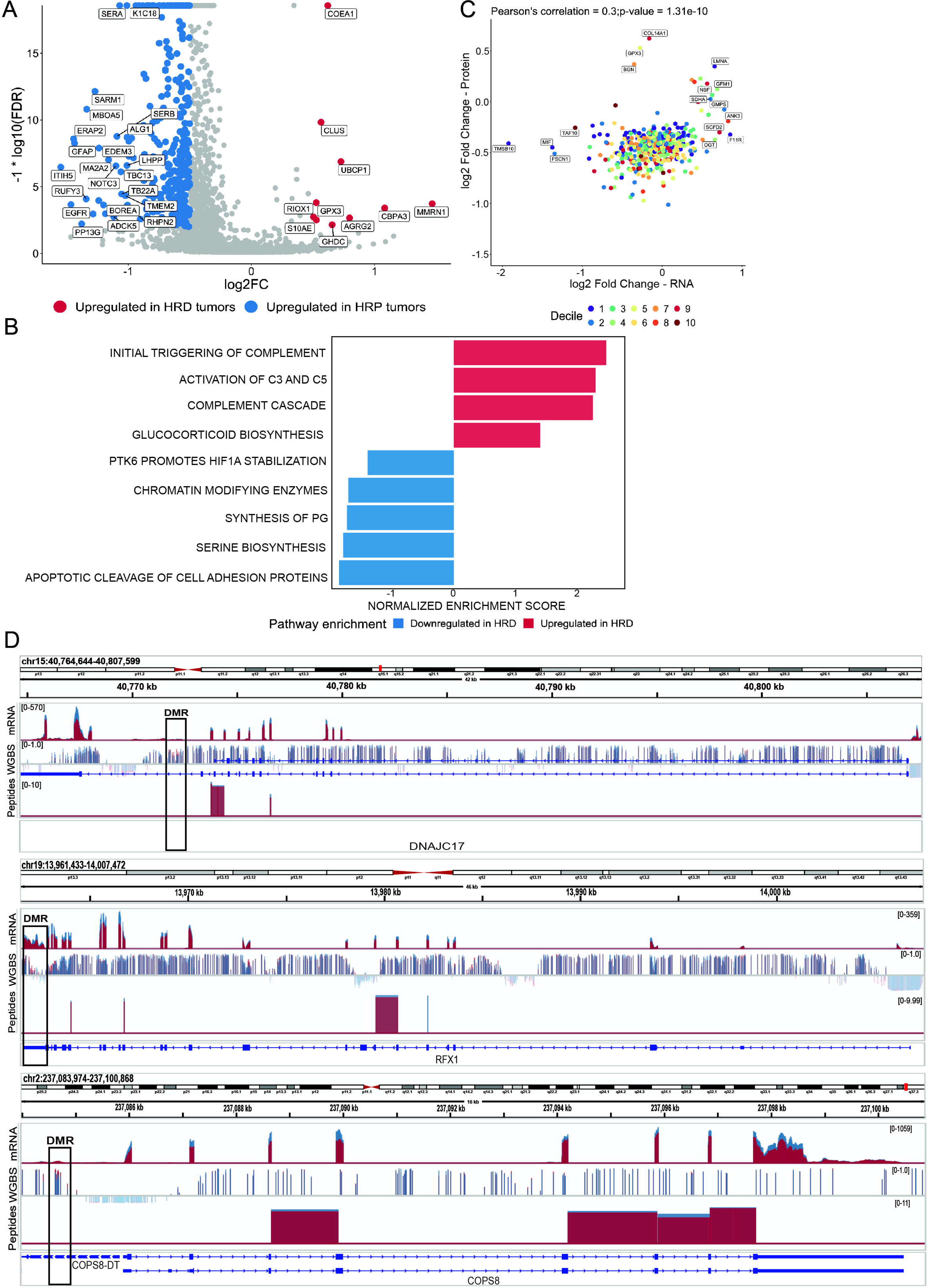
Homologous recombination status reveals shared proteogenomic changes in HGSOC tumors. A. Volcano plot showing significantly upregulated (red) and downregulated (blue) proteins in HRD samples compared to HRP samples (threshold: log2 fold change ± 0.5; FDR < 0.01). B. Gene set enrichment analysis (GSEA) for upregulated (red) and downregulated (blue) proteins in HRD samples (Reactome gene set; p-value < 0.001). C. Comparison of protein and gene log fold changes between HRD and HRP samples. Each gene is colored by protein abundance decile (purple/1= most abundant proteins and dark red/10 = least abundant proteins). D. IGV screenshots of peptide and mRNA gene expression levels for three significantly upregulated proteins/genes in HRP tumors as evidenced by three omics data sets (top - RNA-seq: pvalue < 0.01; middle - differentially methylated regions from WGBS data: pvalue< 0.05 and DistanceTSS < 3000; bottom - proteomics: FDR < 0.05).

Differential expression analysis between primary and recurrent tumors identified 65 proteins differentially expressed between the two groups (1.3% of total proteins, log2FC > +/- 0.5; FDR < 0.01), where 31 proteins were upregulated and 34 were downregulated in primary tumors (**Supp Figure 3A, B; Supp Table 11**). Proteins upregulated in recurrent tumors, like CD5L, CILP, and TINAGL1 have been predicted to be secreted in extracellular space based on the COMPARTMENT database ^44^ and can be further studied as potential biomarkers of chemoresistance for HGSOC. Five upregulated proteins in recurrent tumors in this analysis (TMEM205, PDK3, EPS8, TK1, IGF2BP1) are known to promote chemoresistance in various cancers, indicating shared mechanisms for chemoresistance between ovarian and other cancers ^45–50^. In patients with serial recurrent samples collected throughout disease course, relapse proteins with reduced expression across recurrence events were enriched in pathways associated with tumorigenesis, epithelial mesenchymal transition upon TGF-beta stimulation, and complement coagulation cascade (linear regression analysis; FDR < 0.05; **Supp Table 12**). Pathways associated with oxidative phosphorylation were enriched for proteins with increased expression across recurrence events and have been enriched in other refractory chemoresistant HGSOC tumors in a recent study ^51^. These results indicate that the proteomic landscape of HGSOC recurrent tumors is conserved throughout recurrence and the development of chemo-resistance, as we have previously reported with other genomic data types ^33^. The small number of detected changes between matched primary and recurrent tumors and the pathways identified as altered across recurrent events suggest that primary tumors already had the machinery in place for metastasis and chemoresistance prior to diagnosis and sampling for this study.

### Homologous recombination deficiency drives molecular subtypes identified in HGSOC tumors

Previous studies have identified four transcriptomic HGSOC molecular subtypes, labeled as differentiated, immunoreactive, mesenchymal, and proliferative, based on gene expression data ^52^. The Clinical Proteomic Tumor Analysis Consortium (CPTAC) study using protein abundance data suggested five molecular subtypes from 169 HGSOC samples, four of which corresponded to the transcriptionally identified subtypes (HGSOC-CPTAC) ^22^. Only the mesenchymal subtype was reproducibly identified when a large subset of HGSOC-CPTAC samples (n=103) were re-processed using SWATH-MS ^53^. In our data, Weighted Gene Co-expression Network Analysis (WGCNA) modules derived using proteins from the five molecular subtypes from the HGSOC-CPTAC study correlated significantly with stromal content in our data set compared to other clinical annotations (**Supp Figure 4A**). Our findings here, combined with other studies, suggest that previously described molecular subtypes in HGSOC may be markers of stromal infiltration within tumors rather than cancer cell-specific subtypes ^54,55^.

Consensus clustering of protein expression data in 59 tumor samples identified four stable clusters, which correlated with 5 WGCNA modules (**Supp Figure 4B,C**). Using WGCNA’s module trait correlations, we sought to understand the underlying biology of the samples defined by consensus clustering. The turquoise module was enriched in KEGG pathways like DNA replication, repair, and protein processing in the endoplasmic reticulum and was significantly positively correlated to cluster 3 (r = 0.41, p-value = 0.001). This specific cluster of predominantly HRP tumors (67%) was also negatively correlated to the brown WGCNA module (-0.41; p-value = 0.001), whose co-expressed proteins were associated with immune pathways (complement coagulation cascade, staph aureus infection, neutrophil extracellular trap formation, and natural killer cell-mediated cytotoxicity, **Supp Table 13**). We found a positive correlation between the yellow WGCNA module enriched in metabolic pathways (amino acid, fructose, mannose, butanoate metabolism, and oxidative phosphorylation; **Supp Table 13**) and cluster 4, which comprised mainly of HRD tumors (70%, p-value = 0.025). Patients whose primary tumors were in clusters 1 and 4 had significantly longer survival than those in clusters 2 and 3 (**Supp Figure 4D**). Thus, we identified a subset of HRP tumors whose proteins were associated with DNA replication and repair and negatively associated with immune signaling. We also identified a subgroup of HRD tumors whose proteins were implicated in metabolic pathways. The robust clustering of tumor samples and their correlations with modules of co-expressed proteins were symptomatic of the differences in the biology of HRD and HRP tumors.

### Differences in HGSOC tumor proteome are driven by homologous recombination deficiency status

We identified 334 differentially expressed proteins based on HR status (log2FC +/- 0.5, FDR < 0.05; **Figure 2A**), of which 10 proteins were significantly upregulated and 324 proteins were significantly downregulated in HRD tumors (**Supp Table 14**). Proteins like POLD1, POLD2, PCNA, XRCC1 and MSH6 which are involved in homologous recombination repair, DNA replication, base excision and mismatch repair were downregulated in HRD tumors. PAX8 and WT1, which are known to be overexpressed in ovarian cancers, were significantly upregulated in HRP tumors (FDR < 0.01). EGFR, ERBB2, and MIF, which were significantly upregulated in HRP tumors, (FDR < 0.01) are currently being studied in phase 2 and 3 clinical trials as biomarkers for ovarian, breast and lung cancers in the Early Detection Research Network (EDRN) ^56^. Proteins upregulated in HRD tumors were enriched in complement cascade-related pathways like the initial triggering of complement and activation of c3 and c5 (p-value < 0.001, **Figure 2B**). Downregulated proteins from HRD tumors were enriched in pathways like the chromatin modifying enzymes and serine biosynthesis (**Figure 2B, Supp Table 15**). Three proteins from the serine biosynthesis pathway (PHGDH, PSAT1, and PSPH) are sequentially involved in serine production and were downregulated in HRD tumors, and therefore conversely upregulated in HRP tumors. Upregulation of the serine biosynthesis pathway has been reported in many cancer types and an improved understanding of these pathways can provide translational opportunities for drug development in HGSOC^57^. Differential expression analysis between stromal HRD and HRP tumors revealed that proteins upregulated in the stroma of HRP tumors were enriched in immune related pathways, like type II interferon signaling, graft versus host disease, allograft rejection and CD40 signaling (FDR < 0.05; **Supp Table 16**). Taken together, the findings from discrete analysis of DEPs in tumors versus stroma suggest that HRD tumors have an active immune system within tumor beds while in HRP tumors this immune activity may be localized to the stroma surrounding the tumors. We also performed differential expression analysis between 58 HRD and 45 HRP samples from the HGSOC-CPTAC study with the specific proteomic informatics parameters used in our analysis. Proteins upregulated in HRD tumors were again enriched in immune pathways such as natural killer cell-mediated cytotoxicity, dental caries, and human cytomegalovirus (HCMV) infection (p-value < 0.001; **Supp Table 17**). Combined with our study, these findings highlight the role of homologous recombination status in immune activation and suppression in HGSOC.

### Homologous recombination status reveals shared proteogenomic changes in HGSOC tumors

In a previous study ^33^, we performed whole genome bisulfite sequencing and transcriptome sequencing in primary and recurrent tumors from patients with stage III/IV HGSOC. In this study, we conducted proteomic profiling on the same set of samples (**Figure 1A**, **Supp Table 2**). The data from both studies were then used for proteogenomic analysis. Among the comparisons we examined, analysis of HRD status showed the most potent effect on the HGSOC proteome, hence we used this comparison to simultaneously assess the shared proteomic and transcriptomic changes caused by HRD status. We focused on 470 transcripts and proteins significantly associated with HRD status (protein FDR < 0.001 and quantified in > 80% samples; mRNA FDR < 0.05; 10 overlapping genes/proteins). Changes in protein expression based on HRD status correlated well with changes in gene expression (Pearson’s correlation =0.3, p-value =1.677e-10). Still, we observed a wider range of gene abundances than protein abundances (**Figure 2C**) which has been observed at the single cell level in other studies ^58^. Macrophage migration inhibitory factor (MIF) was upregulated in HRP tumors at both the protein and transcript levels in our data and also in the HGSOC-CPTAC data. It was one of the top 10 most abundant proteins in our dataset and was quantified in all 59 tumor samples (Supp **Table 3**). MIF expression correlates with increased macrophage infiltration, promoting tumor invasiveness and immune escape in ovarian cancers ^59–62^. A four-protein panel that included MIF with CA125 was better able to distinguish ovarian cancer cases from healthy controls ^63^. This is the first report on MIF and its association with homologous recombination deficiency, which is supported by both bulk RNA-seq and proteomic analysis and an independent HGSOC data set ^53^. The many genes with association of HRD status with only mRNA or protein, but not both, might be attributed to biological factors like translation, protein/gene stability, post-translational modifications, and technical factors.

We next compared the results of calling differentially methylated regions between HRD and HRP samples to results from the differential expression analysis on RNA-seq and proteomic data. Twenty-eight differentially methylated regions (DMRs) that were hypomethylated for genes in HRP tumors overlapped with proteins upregulated in the HRP tumors. RFX1, COPS8, and DNAJC17 were upregulated in HRP tumors, as evidenced by all three omics data sets; proteomics, bulk RNA-seq, and WGBS (proteomics: FDR < 0.05, RNA-seq: p-value < 0.01 and DMRs: p-value< 0.05 and DistanceTSS < 3000; **Supp Table 18**; **Figure 2D**). RFX1 and COPS8 have been implicated in tumor growth, metastasis, or EMT in other cancers, while very little is known about DNAJC17 from the perspective of cancer ^64,65^. These are the first results to show an association with all three of these genes and proteins specifically with ovarian cancer and homologous recombination deficiency.

### Increased presence of immune cells and their localization is associated with chemotherapy and HRD status in HGSOC tumors

Proteins and genes upregulated in HRD samples in our study and the HGSOC-CPTAC (Clinical Proteomic Tumor Analysis Consortium) data were enriched in pathways associated with the innate immune system. Thus, we wanted to understand how the tumor’s ability to repair DNA damage drives the immune system toward activation or suppression with the added context of chemotherapy’s influence on the immune system, which this dataset could uniquely interrogate. We used the tool HoverNet, which uses a convolution neural network to estimate the quantity of different cell types, including inflammatory (immune), connective, epithelial, neoplastic, and necrotic cell types in HGSOC tissue sections, which included regions of both the tumor and surrounding stroma (**Figure 3A**). We found that tissues from recurrent tumor sites, irrespective of their HRD status, had a higher number of immune and necrotic cells than tissues from primary tumor sites (**Figure 3B, Supp Fig 5A**). When we focused only on the neoplastic regions of the tumors used for proteomic analysis, we found that tumors with germline *BRCA1/2* deleterious variants had significantly increased inflammatory, connective, and necrotic subtypes and decreased neoplastic cells compared to other tumors without those variants (**Figure 3C**).

**Figure 3:**
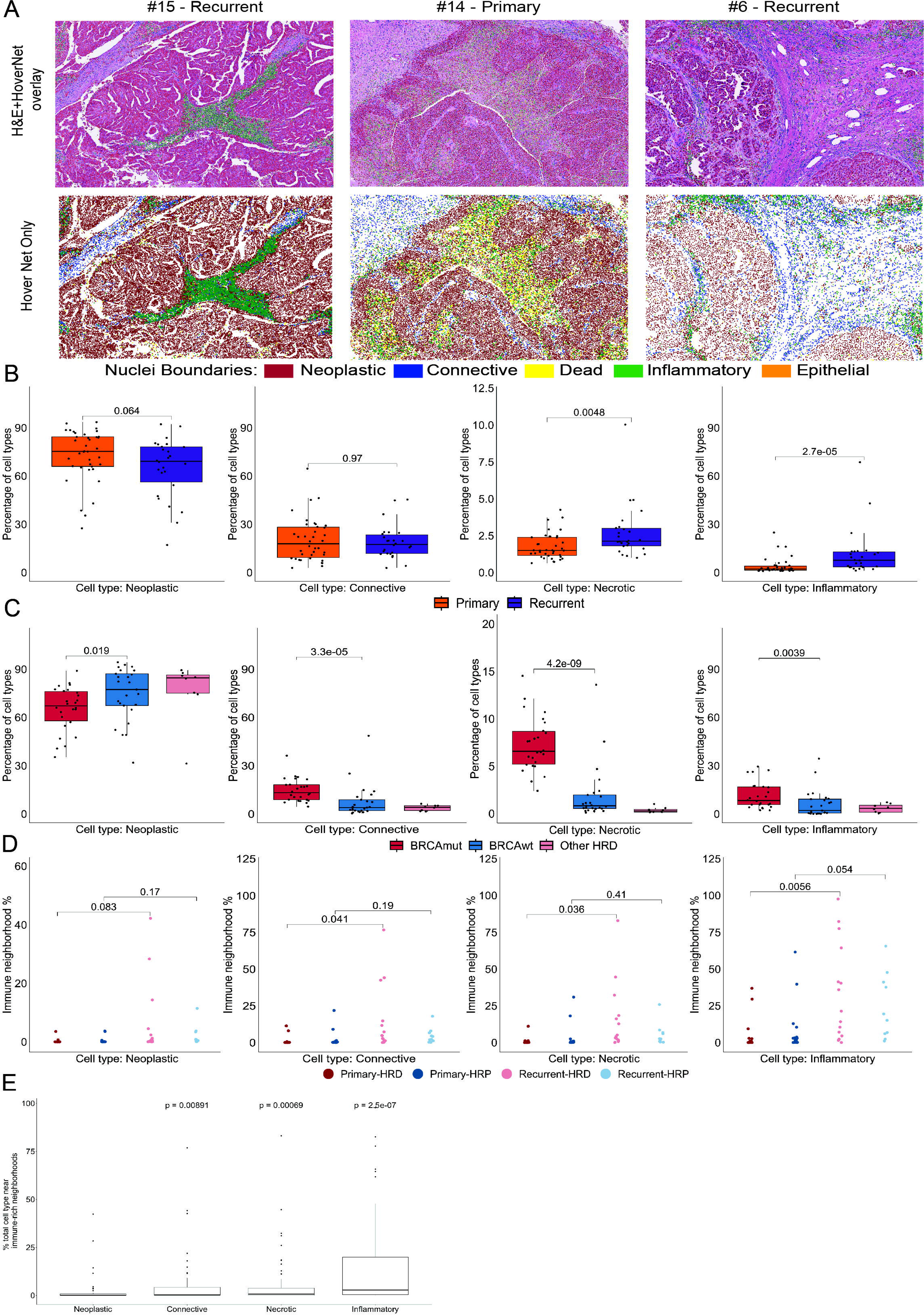
Increased presence of immune cells and their localization is associated with recurrence and HRD status in HGSOC tumors. A. Three representative cropped H&E stained images overlayed with nuclear classification information from HoverNet analysis, which uses convolution neural network to estimate the quantity of different cell types, including inflammatory (immune), connective, epithelial, neoplastic, and dead (necrotic) cell types. HoverNet cell types are colored by bars as indicated below panel. B. Boxplots of percentages of cell types in HGSOC tumors and tumor microenvironments grouped by recurrence status (quantified from HoverNet annotated H&E stained images). C. Boxplots of percentages of cell types in dense neoplastic regions of HGSOC tumors grouped by HRD status (quantified from HoverNet annotated H&E stained images). D. Recurrent tumors have a significantly higher number of immune-rich regions than primary tumors, with localization of immune cells driven by HRD status. E. Inflammatory, necrotic, and connective cell types were significantly closer to immune-rich regions than neoplastic cells, irrespective of recurrence and HRD status.

**Figure 5:**
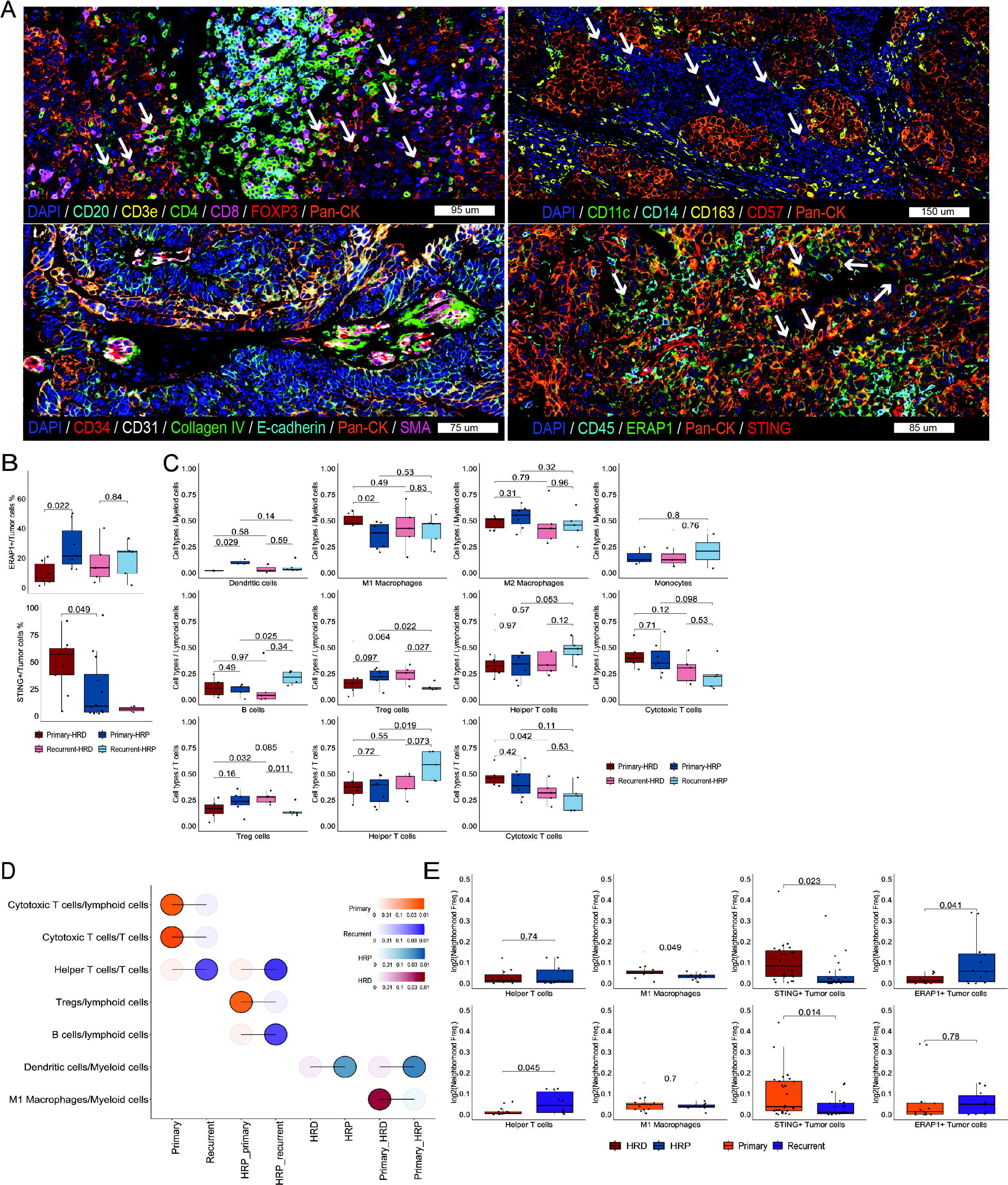
Spatial Analysis of Cellular Changes in HGSOC: Assessing the Impact of HRD Status and Recurrence. A. Four images demonstrating specificity of our Phenocycler imaging. White arrows in the figure point T-regulatory cells (top-left), natural killer cells (top-right), tumor cells positive for ERAP1 and STING expression (bottom-right). Bottom-left shows blood vessels adjacent to tumor tissue. B. (Top) Boxplot showing increased levels of ERAP1+ tumor cells in HRP tumors, particularly HRP primary tumors, compared to HRD primary tumors. (Bottom) Boxplot showing STING+ tumor cells significantly higher in tumors with germline *BRCA1/2* deleterious variants as compared to tumors with no germline *BRCA1/2* deleterious variants and those with somatic *BRCA1/RAD51C* promoter hypermethylation C. Boxplots showing statistically significantly different cell type ratios driven by HRD and recurrence status. D. Bubble plot summarizing significantly different cell type ratios from panel B. E. Boxplots showing statistically significantly different frequencies of neighborhoods driven by HRD and recurrence status.

We performed nearest neighbor analysis to identify HoverNet annotated cell types near immune clusters using images from the whole tissue of HGSOC tumors, including regions of both the tumor and surrounding stroma. We observed that after chemotherapy, only HRD tumors had significantly increased immune clusters near connective and necrotic cells (**Figure 3D**). Overall, connective, immune, and necrotic cells had a significantly higher number of immune cells in their proximity than neoplastic cells (**Figure 3E**). This suggested that inflammatory cells acted in concert with each other based on their proximity to other immune cells and were localized near stromal and necrotic cells, while HGSOC tumor cells had comparatively reduced immune cells in their proximity. Interestingly, tumors from the lymph node, spleen, abdominal wall, omentum, and pelvis had significantly higher immune clusters than tumors from the ovary (**Supp Figure 5B**). When we focused on only tumor specific regions of interest (ROIs) for each tissue, effectively removing surrounding stroma from our analysis, previously identified differences in **Figure 3D** were no longer significant (**Supp Figure 5C**). Overall, this implied that although HRP tumors had an overall increase in immune cell count after chemotherapy, they were more diffuse and not clustered around any specific cell type unlike the cell type specific clustering seen in HRD tumors (**Figure 3D**). The proximity of neoplastic cells with immune clusters was associated with significantly improved survival in HGSOC patients (**Supp Figure 5D**), which was not significant when we calculated the proximities only on tumor specific ROIs for each tissue (**Supp Figure 5E**). The difference in significance after focusing on tumor specific ROIs indicated that, overall, HRD tumors had immune clusters near connective cells and their proximity to neoplastic cells influenced survival in HGSOC patients. Nearest neighbor analysis and total cell type count showed an overall increase in immune cells within HGSOC tissues after chemotherapy. At the same time, the localization of those immune cells was associated with HRD status.

### HRD status is associated with TCR CDR3 repertoires of HGSOC tumor infiltrating T cells and neoantigen count

Given the differences observed in immune cell content based on HRD status, we interrogated the T cell composition and repertoire using TCR-seq on our bulk RNAseq dataset from these tumors. The total number of unique clonotypes (TCR CDR3 sequences) was positively correlated to the number of immune cells within each tumor (p-value = 0.00084, r = 0.46). Clonotype count was also negatively correlated to the number of neoplastic cells (p-value = 0.022, r = -0.33), with no significant correlations observed for connective and necrotic cell types identified using HoverNet (**Figure 4A**). These data demonstrated that the cells annotated by HoverNet imaging analysis as “inflammatory” were immune-related cell types, a subset of which were T cells with distinct CDR3 sequences. To quantify the TCR CDR3 sequence overlap between samples, we calculated the Morisita-Horn index (ranges from 0 indicating no similarity or colocalization to 1 for the two structures being identical or perfectly colocalized) for all sample pairs within HRD and HRP tumors, irrespective of recurrence status (**Figure 4B; left panel**). HRD tumors showed a significantly higher value than HRP tumors, suggesting that HRD tumors have a more shared TCR repertoire compared to HRP tumors, whose repertoire was largely private and restricted to individual tumors. HRD tumors had a significantly higher number of shared clonotypes with other HRD tumors which confirmed this finding (**Figure 4B; right panel**), and this trend has also been observed in an independent HGSOC cohort ^5^ (**Supp Figure 5F**). This was exemplified by increased intrapatient shared clonotype counts in patients with HRD tumors compared to HRP tumors (**Figure 4C**). Patients with an overall higher Gini-Simpson index or clonotype diversity across their primary and recurrent tumors showed significantly improved survival, which has been observed in other cancers ^66^ (**Figure 4D**).

**Figure 4:**
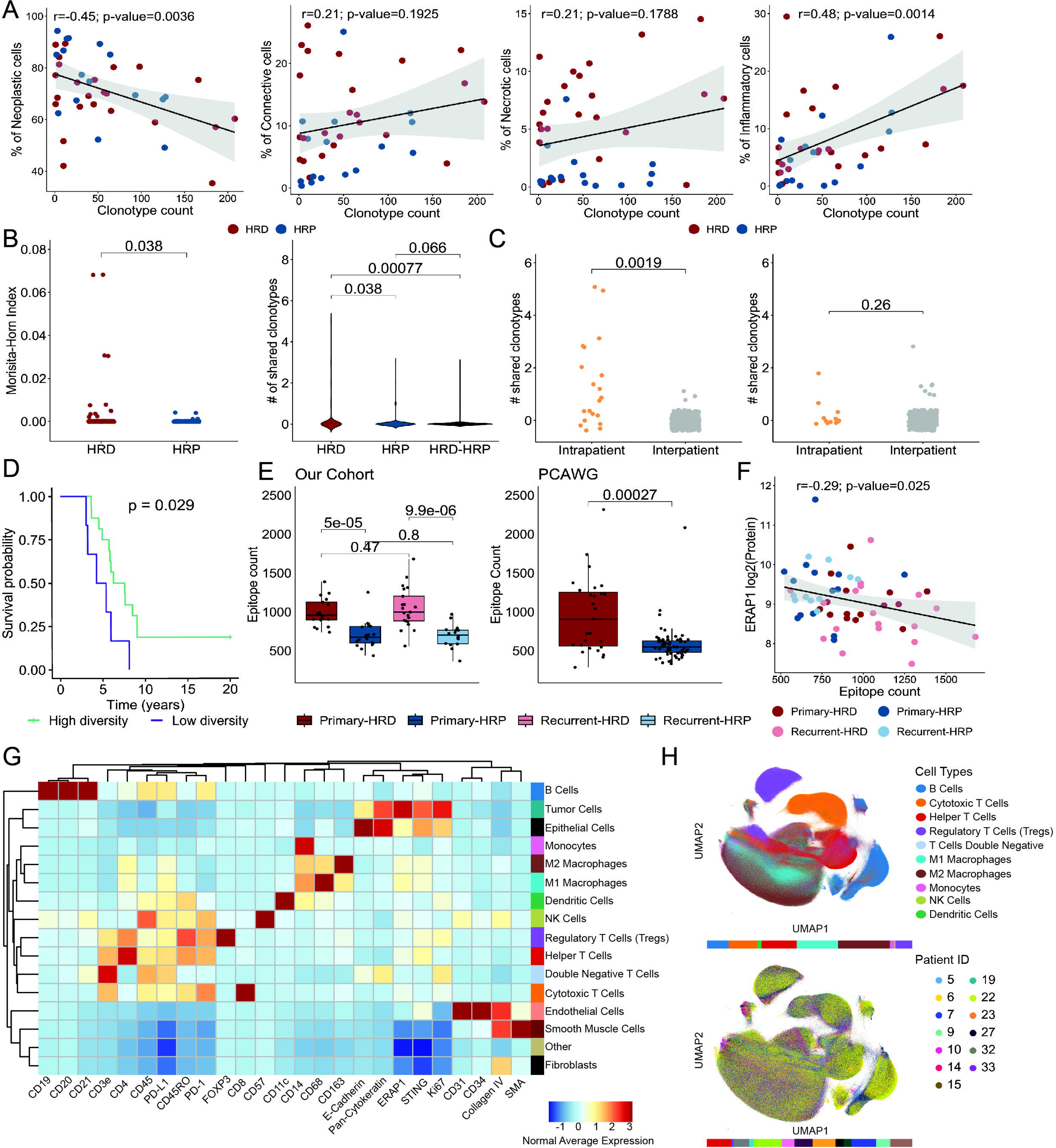
HRD status is associated with TCR CDR3 repertoires of HGSOC tumor infiltrating T cells and neoantigen count. A. Scatter plots showing significant positive and negative correlations between clonotype count (TCR-seq) and the percentage of immune and neoplastic cells quantified from HoverNet analysis. B. (Left) TCR-seq analysis using bulk RNA-seq data shows that HRD tumors have a significantly higher Morisita-Horn index than HRP tumors. (Right) The number of shared clonotypes within and across the two groups of samples. HRD tumors had a significantly higher number of shared clonotypes with other HRD tumors. C. HRD tumors have a significantly higher number of shared clonotypes within patients (Intrapatient count) compared to across different patients - Interpatient count (left-panel) which is not observed in HRP tumors. D. High average Gini-Simpson index or clonotype diversity was associated with improved survival in HGSOC patients (Gini-Simpson index > 0.77 = High clonotype diversity). E. (Left) HRD tumors have a significantly higher number of novel epitopes compared to HRP tumors, irrespective of recurrence status and (Right) that is also reflected in an independent public data set from the Pan-Cancer Analysis of Whole Genomes (PCAWG). F. Scatter plot of ERAP1 protein expression levels vs epitope count shows a significant negative correlation in HGSOC tumors. G. Cell phenotypes in Phenocycler fusion data were defined by unsupervised clustering based on expression of cell lineage and structural markers. H. UMAP representation of all immune cells from Phenocycler fusion data colored by cell type (top-panel) and Patient ID (bottom-panel).

We also found that the number of neoantigens quantified in HRD tumors was significantly higher than in HRP tumors, irrespective of recurrence status (**Figure 4E; left panel**). This was also observed in an independent HGSOC cohort from the Pan-Cancer Analysis of Whole Genomes (PCAWG) study ^5,23^ (**Figure 4E; right panel)**. We did not observe any correlation between TCR clonotype count or diversity (measured as Gini-Simpson index) with novel epitopes. The variability of tumor-infiltrating T cells in recognizing tumor tissue and the presence of bystander T cells may be the reason for this lack of correlation ^67,68^. The intratumoral T cells may also recognize peptide-MHC combinations that were previously present in the tumor but were subsequently lost due to editing of the antigen ^69,70^. ERAP1 and ERAP2 have non-overlapping functions for trimming endogenous cell peptides transported to the endoplasmic reticulum to present to immune cells via MHC class I molecules ^71–74^. ERAP1 protein expression levels were negatively correlated to neoantigens quantified using RNA-seq data (p-value = 0.025, r = -0.3, **Figure 4F**). ERAP1 and ERAP2 were both significantly upregulated in HRP tumors, with ERAP2 falling within the top 5 differentially expressed proteins. Studies have shown over-expression of ERAP1/2 leads to tumor antigen destruction and subsequent immune suppression in colorectal carcinoma and melanoma ^75,76^. ERAP1 should be further studied as a target in combination with immunotherapy, especially for patients with HRP tumors. These results showed that homologous recombination deficiency was a driver of the HGSOC immune landscape affecting TCR repertoires and neoantigens within these tumors.

### Increased Numbers of ERAP1+ Tumor Cells Observed in HRP tumors when Mapping Spatial Proteomic Immune Landscape

We employed a 26-plex antibody panel (**Supp Table 19**) to explore the single-cell spatial relationships of immune cells and homologous recombination deficient/proficient (HRD/P) tumors in paired primary and recurrent HGSOC using Akoya’s multiplex spatial technology ^77^. This study was conducted on 23 tissues collected from various sites, and regions of interest (ROIs) were manually annotated on the H&E images (**Supp Figure 6A**). Cell phenotypes were defined by unsupervised clustering based on expression of cell lineage and structural markers (**Figure 4G**). A total of 36.3 million cells were identified from 23 HGSOC tissue samples and classified into 19 cell subtypes (**Supp Figure 6B, Supp Table 20**). The single-cell spatial maps in **Supp Figure 6C**, accompanied with bar charts below, illustrate the diversity in the cellular composition and spatial organization of the tumor microenvironment of HGSOC tissues in four representative images. The t-SNE projection of cells from paired primary and recurrent tumors of individual patients showed the varying cellular compositions of HGSOC before and after chemotherapy (**Supp Figure 7**). Generally, cellular compositions diversified in recurrences, but only slightly. UMAP projections of immune cells in **Figure 4H** indicated varying proportions of 10 different immune cell populations, with cells colored by patient ID shown in the bottom panel. **Figure 5A** displays examples of prominent cell types that were identified. White arrows in the figure indicate Tregs (Top-left), NK cells (Top-right), tumor cells positive for ERAP1 and STING expression (Bottom-right). M2 macrophages were the most abundant immune cell type (25% of total immune cells) followed by M1 macrophages (20% of total immune cells), a finding also observed in ovarian cancer samples from The Cancer Immunome Atlas ^78^. Percentages of different cell types identified by CODEX and HoverNet (**Figure 4A**) were significantly positively correlated for the 23 HGSOC tissues (**Supp Figure 8A,B**). Notably, M1 and M2 macrophage counts from Akoya’s data were poorly correlated with inflammatory cell percentages from HoverNet (**Supp Figure 8A,B**) indicating HoverNet’s limitations in detecting specific types of immune cells.

**Figure 6:**
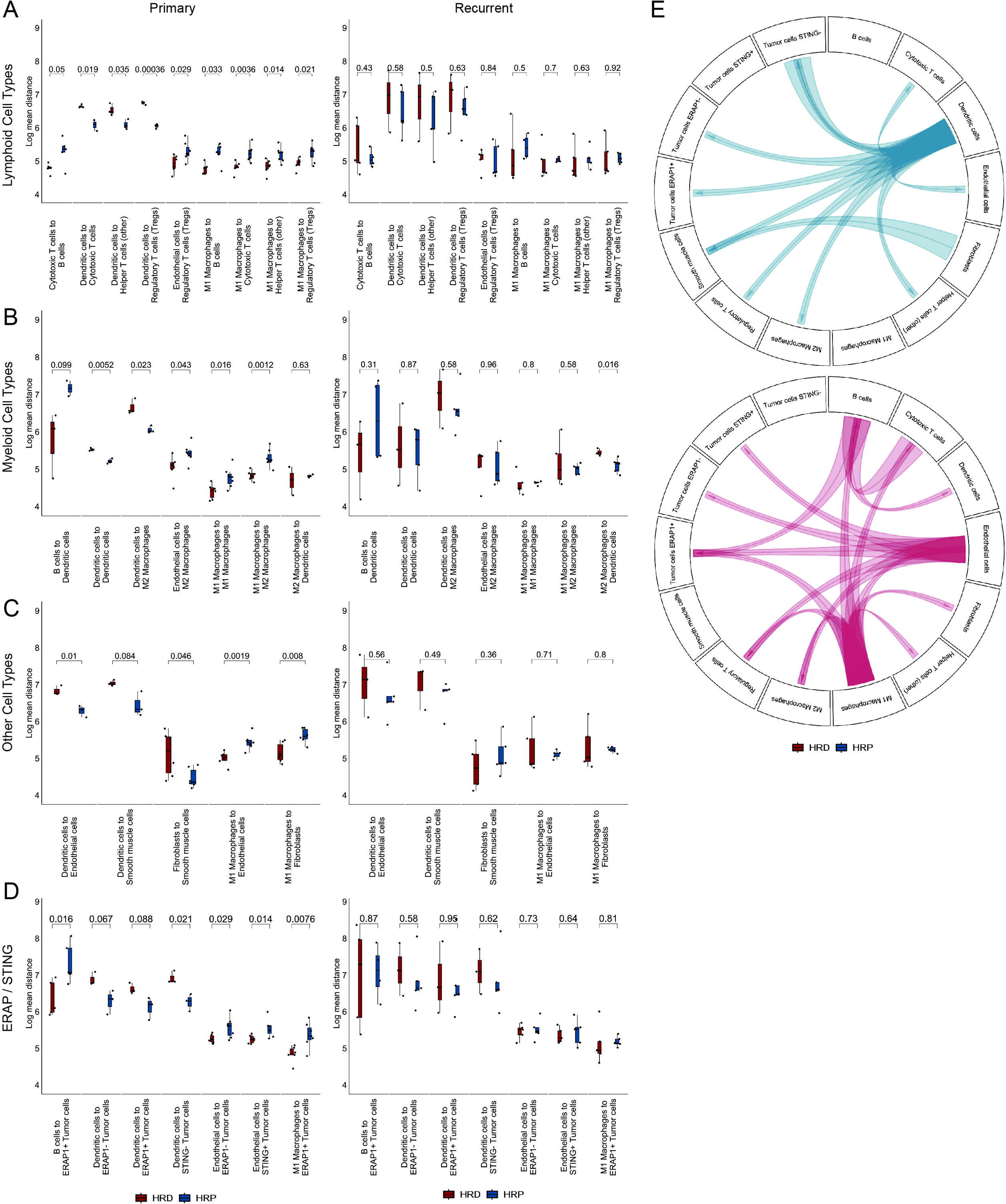
Spatial Proximity Analysis Reveals Significant Distances between HRD and HRP Primary Tumors. A-D. Spatial proximity analysis outlining the localization of cell types relative to each other reveals a larger number of significant differences between the HRD and HRP groups in the primary tumors. Panels are groups by cell type: Lymphoid, Myeloid, Other, and ERAP/STING are represented in A-D, respectively. E. Circos plots highlighting statistically significant proximity analysis results from A-D for primary tumors.

We observed increased levels of ERAP1+ tumor cells in HRP tumors, particularly HRP primary tumors when compared to HRD primary tumors (p-value = 0.022; **Figure 5B, top panel**). These results supported our findings from the differential protein expression analysis where we found ERAP1 and ERAP2 levels significantly upregulated in HRP tumors compared to HRD tumors (**Figure 2A**). Studies have shown STING to be upregulated in ovarian tumors and cell lines with BRCA1/2 mutations ^12,13^. Similarly in our study, STING+ tumor cells were significantly higher in tumors with germline *BRCA1/2* deleterious variants compared to tumors with no germline *BRCA1/2* deleterious variants and those with somatic *BRCA1/RAD51C* promoter hypermethylation (p-value = 0.049; **Figure 5B, bottom panel**).

Treg levels changed inversely after chemotherapy, based on HRD status; in HRD tumors Tregs increased, while in HRP tumors Tregs decreased post-chemotherapy (**Figure 5C,D)**. The increases of Tregs in HRD recurrent tumors may be related to upregulation of TK1, one of the differentially expressed proteins found in our cohort. Tregs have been shown to modulate the expression of TK1 which can in turn affect progression and occurrence of tumors ^79^.

Consistent with previous observations from an image analysis of HGSOC ^80^, CD11c+ dendritic cells (DCs) were significantly higher in HRP tumors compared to HRD tumors (**Figure 5C,D)**. Although higher DC counts typically indicate improved immune surveillance ^81–83^, it is possible these cells are trapped within the stroma as described in Papadas et al ^84^ and unable to penetrate the tumors indicating a subdued immune response.

We saw a significant increase in M1 macrophage count in HRD tumors compared to HRP tumors (**Figure 5C,D)**. For tissues exposed to chemotherapy, HRD status did not cause any statistically significant differences in cell counts. In other breast and ovarian studies, increase in M1 macrophage infiltration and M1-like macrophage genes were associated with higher mutation rates ^85^, longer survival ^86^ and higher HRD scores ^87^. This result confirms our Hovernet results in **Figure 3C** showing significantly increased inflammatory subtypes in HRD tumors.

Next, we performed nearest neighbor analysis to identify common cellular neighborhoods across HGSOC tissues, driven by either recurrence or homologous recombination deficiency status (**Supp Figure 8C,D, E**). Our study revealed that HRD tumors have a significantly higher presence of cells in the neighborhoods of M1 macrophages compared to HRP tumors (**Figure 5E**). Additionally, recurrent tumors exhibited a significantly greater cell count in the neighborhoods of T helper cells when contrasted with primary tumors (**Figure 5E**). We observed significantly higher number of cells near neighborhoods of ERAP1+ tumor cells in HRP tumors, and significantly higher number of cells near neighborhoods of STING+ tumor cells in HRD tumors (**Figure 5E**). The localization patterns observed in the nearest neighbor analysis mirrors the cellular abundance shifts between different groups of samples described above.

To further explore the function of immune cells as inferred from their vicinity to other cells, we performed spatial proximity analysis. Based on HRD status, we found more significant differences within primary tumors compared to recurrent tumors, where only the distance of M2 macrophages to DCs were significantly closer in HRP compared to HRD recurrent tumors (**Figure 6A-E**). In general, DCs were significantly closer to various immune cells (Helper T cells, Cytotoxic T cells, regulatory T cells, M2 macrophages) and other DCs in HRP tumors compared to HRD tumors. In contrast, DCs were significantly closer to only B cells in HRD tumors compared to HRP tumors (**Figure 6A-E**). These two observations again indicate DCs, as well as other immune cells, may be trapped in the stromal compartment, unable to penetrate the tumor bed.

In HRD tumors, M1 macrophages were significantly closer to most immune cell types (cytotoxic T cells, B cells, M2 macrophages, regulatory T cells, Helper T cells, other M1 macrophages) and ERAP1+ tumor cells compared to HRP tumors (**Figure 6A-E**). STING has been shown to promote neoangiogenesis by increasing levels of VEGF-A in *BRCA1* mutant tumors ^12^. Preliminary evidence from 222 cases of paired primary and recurrent HGSOC from 111 patients showed that somatic *BRCA* mutated samples had increased VEGFA levels ^16^. In our study, STING+ tumor cells were significantly closer to endothelial cells in HRD tumors compared to HRP tumors while also being significantly more abundant in a subset of HRD tumors with germline *BRCA1/2* deleterious variants.

Cytotoxic T cells were significantly closer to M1 macrophages and to B cells in HRD tumors compared to HRP tumors. A greater proximity between cytotoxic T cells, B cells and M1 macrophages in HRD tumors might be indicative of a more coordinated and effective immune response, as also evidenced in a previous study of *BRCA1/2* mutated HGSOC ^80^ (**Figure 6A-E**). Overall, our results indicate higher involvement of immune cell populations within HRD compared to HRP based on cellular location within tumors. The immune environment and tumor characteristics have been previously shown to vary based on the mutation status as well as the tumor’s anatomical site ^88^. These results underscore the importance of understanding not only the genetic and epigenetic profiles of a tumor, but also the spatial topology, especially due to the sometimes paradoxical nature of immune cell populations and their core functions in cancer.

## Methods

### Study design

The Women’s Cancer Program Biorepository at Cedars-Sinai Medical Center has collected extensive clinical data and specimens from more than 16,000 patients over 25 years. For this study, we used a set of 32 high-grade serous ovarian cancer (HGSOC) patients diagnosed with Stage III/IV disease for whom paired tumor and adjacent stromal samples were profiled (IRB #0901 and #41922). These cases have extensive clinical history available, including information on cycles and dosage of chemotherapy (**Supp Table 1**). At the time of surgical debulking, germline DNA was collected from circulating blood before chemotherapy, and tumor samples were fresh frozen shortly after collection.

### Proteomics sample processing and data acquisition Protein extraction

Samples were processed as described previously ^89^. Briefly, fresh frozen tumors were embedded in an optimal cutting temperature compound, bisected, and mounted before slides were made for hematoxylin and eosin staining. A single pathologist reviewed all slides to identify regions of epithelial carcinoma. A 3mm core of approximately 50mg was then extracted from a part of pure epithelial carcinoma from the intact frozen tumor, with care not to include any surrounding stroma. These ‘pure tumor’ tissue cores were used for subsequent proteomic analyses. We also extracted regions adjacent to the ‘pure tumor’ tissues which were either a mix of tumor and stromal samples (labeled as 50% stromal content) or stromal samples (labeled as 100% stromal content). Frozen tumors and adjacent stromal samples were thawed on ice, 200 – 400 ml lysis buffer (6M Urea + 0.1% Rapigest) was added, and tissues were homogenized with a polytron. Proteins were extracted by high-pressure barocycling. Protein concentration was assayed using the Pierce BCA assay.

### Data acquisition

Data acquisition was performed as described previously. In short, 4 ml of the digested sample was injected directly into a 200 cm micropillar array column (uPAC, Pharmafluidics) and separated over 120 minutes reversed-phase gradient at 1200 nL/min and 60 C. The eluting peptides were electro-sprayed through a 30 um bore stainless steel emitter (EvoSep) and analyzed on an Orbitrap Lumos using data-independent acquisition (DIA) spanning the 400-1000 m/z range. After analysis of the full m/z range (40 DIA scans), a precursor scan was acquired over the 400-1000 m/z range at 60,000 resolution.

### Peptide assay library

To construct a comprehensive peptide ion library for the analysis of human ovarian cancer, we combined several datasets, both internally generated and from external publicly available resources. First, we utilized a publicly available HGSOC proteomics experiment by downloading raw files from the online data repository and searching them through our internal pipeline for data-dependent acquisition MS analysis as described in Parker et al. ^90^ and performed by others on the same publicly available dataset ^53^. Database searches for both the internal and downloaded external datasets utilized human protein sequences defined in a FASTA database of Swiss-Prot-reviewed, Human canonical reviewed proteome containing 20,406 protein sequences that were downloaded July 2019 and appended with Biognosys indexed retention time (iRT) peptide sequence (Biognosys, Schlieren, Switzerland) and randomized decoy sequences appended. A final, combined consensus spectral library containing all peptide identifications made between the internal and external datasets was compiled. Decoy sequences were appended using the transproteomic pipeline with Spectrast library generation and conversion to TraML format as described previously ^91^. The final library has been provided as a supplemental file to this report. Peptide identification was performed as previously described ^91,92^. MS data and supplementary analysis files were submitted to ProteomeXchange via the PRIDE database under the identifiers PXD023012 for the dilution series, PXD022996 for the small pilot data set, and PXD023040 for the patient tumor proteomic data set.

### RNA-seq library preparation and sequencing

Samples for RNA-seq were processed as described previously ^33^. Briefly, RNA was extracted using the protocol published online by the Prostate Cancer Biorepository network (SOP#006). Sample concentration was measured using the Qubit RNA Broad Range kit and sample quality was measured using the Agilent BioAnalyzer 2100. Sequencing libraries were prepared by adding 1ug of RNA to the TruSeq Stranded Total RNA Kit with Ribo- and Mito- depletion following the TruSeq standard protocol with 15 cycles of PCR. Libraries were quantified using Qubit RNA Broad Range kit and pooled before being run on one lane of a HiSeq2000 to collect ∼ 1 M reads per library for quality control. PCR duplication rate was estimated in this low coverage sequencing run in 150 bp paired end mode, and library complexity was estimated using PreSeq. Based on the complexity measured in this low coverage sequencing experiment we estimated the maximal coverage that would continue to provide informative measurement of transcripts in the library was ∼ 350 M reads. Each library was then pooled and this pool was sequenced in 2 × 150 bp mode on an Illumina Novaseq 6000, and we generated ∼ 335 million reads from each library. Our data analysis workflow for RNA-Seq has been developed specifically to improve gene feature identification and measurement in archived frozen tissue samples, which can perform poorly using standardized workflows. **RNA-Seq quantification and statistical analysis**

RNA-seq quantification and statistical analysis were performed as described previously^33^. Reads within each fastq file were first trimmed using TrimGalore to remove low quality bases and sample barcodes, retaining reads 75 bp or longer. Each transcriptome was then aligned to hg38 and the Gencodev29 primary assembly ^93^. Genes were quantified with RSEM ^94^ and Kallisto ^95^. Sample-specific gene models were generated using alignments produced with STAR two pass mapping and Stringtie ^96^. Gene expression values were shown as normalized variance stabilizing transformation (vst) counts. To measure RNA abundance, we first obtained BAM alignment quality metrics using Picard (http://broadinstitute.github.io/picard). Samples with less than 90 percent of reads mapped to the correct strand of the reference genome (PCT_CORRECT_STRAND_READS) were omitted. Patients whose primary and recurrent tumors both passed this quality control were retained (n = 50). Read counts were quantified using the R package Salmon ^97^ at transcript level and reads were mapped to Genecode Release 29 (GRCh38) comprehensive gene annotations by R package ‘tximport’ ^98^. To filter out potential artifacts and very low expressed transcripts we retained transcripts with length greater than 300 bp, TPM (Transcripts Per Million) value greater than 0.05 and isoform percentage greater than 1%. Transcripts in blacklist regions ^99^ were also filtered out. We retained transcripts expressed in more than 5 samples, which resulted in 91,411 transcripts from 33,969 genes. Tumor purity was estimated by the degree of heterogeneity of the tumor microenvironment. We applied the R package ‘consensusTME’ ^100^ to estimate cell type specific enrichment scores based on TCGA ovarian cancer data.

### Whole genome bisulfite sequencing (WGBS)

Our workflow for WGBS required a minimum 300ng of high quality gDNA, which was sheared to approximately 175-200 bp using a Covaris sonicator, and bisulfite converted using the EZ DNA Methylation-Lightning Kit (Zymo). Libraries were constructed using the Accel-NGS Methyl-Seq DNA Library Kit (Swift Biosciences, MI), and amplified using no more than 6 cycles of PCR. Libraries were sequenced to at least 30x coverage (on average each base is sequenced thirty times) on the Illumina HiSeq4000 in 150bp paired end mode. This approach generated approximately 400 million read pairs per library, with a bisulfite conversion rate greater than 99%.

### WGBS data processing

WGBS reads were aligned to the human reference genome (build GRCh38) using BISCUIT ^101^. Duplicate reads were marked using Picard Tools ^102^. Methylation rates were called using BISCUIT. CpGs with fewer than 5 reads of coverage were excluded from further analysis. Adapter sequences were trimmed using TrimGalore ^103^, using default parameters for Illumina sequencing platforms. Quality control was performed using PicardTools as well as MultiQC ^104^. Bisulfite non-conversion was checked using the Biscuit QC module in MultiQC. Principal Component (PC) analysis was performed on CpGs with coverage ≥10 and the top 10,000 most variable CpGs were included in the identification of the top 10 PCs using the prcomp function from the stats package in R ^105^.

### Outlier fragment filtration and sample filtration strategies

Data sets used for specific figures are tabulated with the filtration strategy in **Supp Table 21**. We used the mapDIA tool to remove outlier fragments before peptide and protein quantification with MSstats as described in a previous publication ^89^(Data set: SG1). Samples were run in four different mass spectrometry batches. We pooled protein lysates from all samples to generate a ‘pooled’ lysate to run with the four mass spectrometry batches. Samples were randomly assigned to the four batches for a robust study design. BPCA imputation was performed at the protein level (Data set: SG2) on all samples, including pooled samples. Hierarchical clustering of tumor and stromal samples after BPCA imputation revealed outliers with a high percentage of missing values across all quantified proteins. Outliers identified by hierarchical clustering were removed from the downstream analysis, which resulted in the removal of 2 tumors and 1 stromal sample and their replicates(Data set: SG3). One technical replicate per sample with the least missing values at the fragment level was retained for downstream analysis. The resulting set of samples and their fragments with no technical replicates and outliers were re-processed through the mapDIA and MSstats pipeline to remove any outlier fragments further, as described before. The final data set of 96 samples from 32 patients included 60 tumor samples and 56 stromal samples (Data set: SG4).

### Sample re-classification using the stromal score

A recent study of gene expression data has shown that cellular admixtures affect the prognostic capability and reproducibility of ovarian cancer gene signatures and molecular subtypes ^34^. We applied a histology guided micro-dissection method to collect tumor samples that were highly enriched for epithelial cells to prevent stromal contamination of tumor samples. We extended this to include cell annotation and deconvolution methods to prevent stroma contamination from confounding our analyses. We used two separate tools to determine an average stromal score for each sample and re-classified samples as tumor (stromal score < 0.35), stroma (stromal score > 0.55), or a mixture of both (stromal score >0.35 & < 0.55) (**Supp Figure 1A,1C – top panel**). We performed deconvolution analysis using CAM3 with data set SG5 as input (https://github.com/ChiungTingWu/CAM3). We used raw protein intensity values as input for CAM3 and performed deconvolution analysis with default settings and k=2. The random seed generator was set to 4242 for reproducibility. Cluster proportions generated for clusters 1 and 2 were incorporated into the stromal score calculation. H&E stained images in .svs format from profiled tissues were used as input for HoverNet analysis ^106^. A pre-trained model on the PanNuke data set provided by the developers of HoverNet was used for image analysis ^106,107^. The resulting .json file was further analyzed with a custom python script to generate cell type proportions within the regions of tissues sampled for proteomic analysis. A summary stromal score was created using the cell type proportions of neoplastic and connective cell types converted to percentages and then scaled from 0 to 1 for each sample to integrate with CAM cluster proportions. **Supp Figure 1C** - bottom panel showed the changes in sample grouping before and after reclassification, where significant re-distribution occurred for samples previously labeled as “mix.”

### Methods for assaying proteomic landscape of HGSOC

Median log2 protein abundance for 5036 proteins ignoring missing values was calculated across all samples and ranked from highest to lowest mean abundance (Data set: SG6). We calculated the standard deviation of proteins across tumor samples (Data set SG8) and the mean log2 intensity of those proteins. We calculated the mean log2 intensity of proteins across all tumor samples as classified using our stromal score and the number of proteins quantified across the 56 tumor samples (ignoring missing values; Data set SG7). The mean log2 protein intensities were divided into deciles, with the highest abundant proteins in decile 1 and the lowest in decile 10. All pathway analysis was performed using the R package ClusterProfiler. We calculated Pearson’s correlation coefficient for all samples, including replicates for all 5036 proteins using the corrplot package (Data set: SG3) and performed hierarchical clustering of Pearson’s correlation coefficients. For the average replicate correlation coefficient, we first calculated Pearson’s correlation coefficient for every sample replicate with itself, and all other sample replicates. Next, we converted the coefficients to fisher Z-scores and calculated the mean of these scores for all replicates of every sample. Finally, we converted the fisher scores back to Pearson’s correlation coefficients to get the average Pearson’s correlation coefficients within replicates of samples. We used the FisherZ and FisherZInv functions from the DescTools package in R.

### Hierarchical clustering and PCA

We performed principal component analysis using data set SG9 with base R function prcomp on a transposed matrix with samples as rows and proteins as columns. We performed hierarchical clustering using data set SG10 with the ComplexHeatmap package in R. For calculating the Euclidean distance between primary and recurrent tumors; we used only those patient samples whose primary tumor had one or more paired recurrent tumors profiled. The distance between the samples was calculated using the dist2 function in ComplexHeatmap. For primary tumors with more than one paired recurrent tumor, the average distance from primary to all corresponding recurrent tumors from the same patient was used (intra-patient distance). For the inter-patient distance, the average distance of the primary tumor from one patient to all recurrent tumors from all other patients was used.

### Molecular subtypes of HGSOC and incorporation of CPTAC-WGCNA modules

We downloaded the proteins and their correlations to WGCNA-defined modules from the HGSOC-CPTAC study ^22^. We next filtered for proteins with a Pearson’s correlation coefficient of more than 0.5 for each module. If a protein was associated with more than one module, we kept the protein module association with the highest correlation coefficient. We used protein intensities of the 508 identified proteins across 59 samples as input to WGCNA with default parameters (random seed = 4242 and power of 12) which resulted in four modules. We correlated these modules to clinical annotations like recurrence and HRD/P status, along with the stromal score. We performed KEGG enrichment analysis using the ClusterProfiler package and compared the pathway enrichment between different modules using the CompareCluster function.

We performed clustering analysis on the SG10 data set using the ConsensusClustering^108^ tool in R with default parameters (K = 2-10 and random seed= 4242). We next used WGCNA to determine the coexpression of proteins across samples in the SG10 data set, the same data set used for clustering (random seed= 4242 and power = 10). WGCNA-based modules were correlated to clusters, and stromal content of samples and pathway enrichment was done using the ClusterProfiler package described above for CPTAC modules. We performed the survival analysis using packages ‘survminer’ and ‘survival’ in R (https://github.com/kassambara/survminer, https://github.com/doehm/survivoR).

### Differential expression analysis and ANOVA

ANOVA: We performed a two-way ANOVA using the ‘aov()’ base function in R, which does not assume equal variance between groups. ANOVA tests were performed for samples grouped by recurrence status and HRD status. Negative log10 p-values from both tests were plotted as x and y axes with points colored by density using the ‘ggpointdensity’ package in R (https://github.com/LKremer/ggpointdensity).

Differential expression analysis based on HRD status and recurrence for proteomics data was performed using mapDIA on log2 transformed fragment intensities. mapDIA input parameter files are provided in the project github repository. We performed pathway enrichment analysis using ClusterProfiler using the Reactome-gene set downloaded from the human MSigDB collection. DEA for RNA-seq data was performed using a t-test with p-values calculated using Mann Whitney U-test and corrected for multiple testing.

### Differentially Methylated Region - Differentially Expressed Gene Analysis

The Bioconductor package dmrseq ^109^ was utilized to identify DMRs between HRD and HRP samples using default settings. Correlation between DMRs and gene expression was performed by comparing samples with matched WGBS and RNA-seq available for both primary and recurrent tumors. Using GENCODE42, DMRs regions >2kb from any TSS were annotated as “distal” and regions <2kb from TSS were annotated as “promoter”. Distal regions were mapped to the closest genes (10 upstream and 10 downstream) and the promoter to the closest gene and the correlation between their expression and methylation measured, where average beta value of the DMR correlates (using Spearman test) to a change (positive or negative) in expression of the nearby genes.

### Linear regression analysis for serial recurrences

We performed linear regression analysis accounting for stromal content, patient IDs and HRD status to identify changes in protein expression profiles across multiple recurrences (Formula: ∼ stromal score + recurNumber + recurNumber:PatientID + recurNumber:HRD status). We applied weights to coefficients in order to account for the imbalance in samples per group. Weights were calculated based on the ratio of samples in each recurrence group. We performed pathway enrichment analysis using ClusterProfiler using the C2- curated gene set downloaded from the human MSigDB collection.

### TCR-seq analysis from bulk RNA-seq

We used the BAMs from RNA-seq data for 50 HGSOC samples from our previous study (hg38 aligned) ^33^ and downloaded BAMs from RNA-seq data for 74 neoadjuvant HGSOC tumors (hg19 aligned) ^5^. Both data sets were processed using TRUST4 (TCR Receptor Utilities for Solid Tissue) to infer complementarity determining region 3 (CDR3) sequences of tumor-infiltrating T cells in the HGSOC samples ^110^. We used Immunarch^111^ to process TRUST4 results and quantify the Morisita-Horn index and the number of unique CDR3 clonotypes per sample. We used R packages ‘survminer’ and ‘survival’ for survival analysis (https://github.com/kassambara/survminer, https://github.com/doehm/survivoR). For the survival analysis, average gini-Simpson index was calculated across primary and recurrent tumors for every patient. Patients with a gini-Simpson index lower than 0.78 (< second quartile) were labeled as having low clonotype diversity.

### Nearest neighbor analysis; HoverNet data

We performed nearest neighbor analysis to quantify the immune-rich areas within HGSOC tumors and tumor microenvironments using HoverNet annotated nuclei from H&E stained images. Immune-rich regions were defined as regions with greater than 40% immune cells within circular regions of radius 820 pixel in H&E stained images with K=50 nearest neighbors. This allowed us to quantify the number of different cell types with more than 40% immune cells within circular regions of radius 820 pixel around them and determined which cell type was in close proximity to immune cells for every sample. HoverNet annotations overlaid onto H&E stained images were generated using custom python scripts (github link for paper code). We used R packages ‘survminer’ and ‘survival’ for survival analysis (https://github.com/kassambara/survminer, https://github.com/doehm/survivoR). For the survival analysis, average neoplastic cells within immune-rich regions were calculated across primary and recurrent tumors for every patient. If the sample had more than 1% of neoplastic cells near immune-rich regions (> third quartile), they were considered as tissues with a high number of neoplastic cells near immune-rich regions.

### Visualization of mean peptide, mRNA expression levels in igv

RNA-seq: RNA-seq bigwigs were first converted to bedgraph files using ucsc-bigwigtobedgraph function and relevant chromosome were filtered out and saved as separate bedgraph files. Bedtools ‘unionbedg’ function and custom awk script were used to calculate the mean of all samples within HRD and HRP groups and saved as two bedgraph files. The bedgraph files were converted to bigwigs using ucsc-bedgraphtobigwigs function and viewed on igv.

Peptides: Genomic locations of each peptide obtained using PoGo (ref) were added onto peptide expression level matrix. Mean expression levels of all peptides within HRD and HRP groups were saved as two separate bedgraph files. Sorted bedgraph files were converted to bigwigs using ucsc-bedgraphtobigwigs function and viewed on igv.

### Neoantigen Prediction

Briefly, a VCF containing somatic variants identified by Strelka2 ^112^ was annotated by VEP ^113^ with the options --format vcf --vcf --symbol --terms SO --tsl --hgvs --fasta –offline --cache --plugin Frameshift --plugin Wildtype. Expression data from matched RNAseq was added using vcf-expression-annotator ^114^ Resulting VCF was processed using pvacseq version 3.1.1-slim ^115^. The epitope prediction algorithms used were MHCflurry, MHCnuggetsI, MHCnuggetsII, NNalign and NetMHC.

### Homologous recombination status prediction

Homologous recombination status was determined using the Classifier of HOmologous Recombination Deficiency (CHORD) score ^116^. SNV and INDEL information was extracted from Mutect2 VCF files. SV information was extracted from GRIPSS VCF files, the filtered high-confidence output from GRIDSS.

### Immune cell mcIF

FFPE tissue sections of a subset of patients with paired primary and recurrent tumors were selected for mcIF staining using a 26-plex antibody panel on PhenoCycler^®^-Fusion (**Supp Figure 5A**). The 26-plex antibody panel included markers against various immune cell types, immune checkpoint inhibitors, epithelial and stroma-specific markers, and custom markers for validating proteomic differential expression analysis results (**Supp Table 19**).

### Segmentation

Data Analysis was performed using Akoya’s internal software, <u>M</u>ultiplexed Image <u>A</u>nalysis (MIA), which combines multiple existing analysis tools with a Graphical User Interface. Quality control of the data was performed qualitatively on each individual marker image and filtering was conducted to exclude markers and tissue regions with inadequate quality. Nuclear segmentation was first performed using StarDist ^117,118^ method applied to the DAPI channel. Cytoplasm segmentation was then estimated from nuclear expansion by morphological dilation and the centroid of each cell was defined by the x-y coordinates in the image.

### Unsupervised clustering and phenotyping

The mean fluorescent intensity (MFI) of each marker was calculated for each cell from the corresponding expression compartment, e.g., nuclear or cytoplasmic surface, to produce a raw expression table where each row represents a cell and columns are their x-y coordinates and protein MFI expressions. Then z-score normalization was applied across all cells for each marker such that each marker has a mean equal to 0 and a standard deviation equals to 1. Only QC-passed lineage markers were used for downstream analysis. Cells without any marker expression or with extremely large nuclear masks were filtered out from the downstream analysis. After cell segmentation and filtering, there were 36.3 million cells across 23 samples.

The normalized MFI expression table for all markers (columns) and all cells (rows) from each QPTiff image were the input for unsupervised clustering. The Leiden ^119^ algorithm and GPU-accelerated ^120,121^ method was used for clustering. For Leiden clusters, resolution equals to 1-6 were tested and the optimal resolution = 1 was chosen for manual annotations to assign cell phenotypes based on marker expression pattern in hierarchical clustering heatmap and their location on the images. Clusters with similar expression profiles were combined into one phenotype and a new heatmap with re-averaged protein expressions and cell phenotypes was generated. For tumor cells classified as ERAP1+/- and STING+/-, tumor cell populations were clustered using K means clustering, with 5 clusters and then the tumor cells in top 3 clusters were assigned as ERAP1+ or STING+ while the rest were labeled as ERAP- or STING-.

### UMAP/t-SNE

Dimensionality reduction was applied for data visualization by means of tSNE using Scanpy ^119^ and Rapids ^120,121^ implementations. tSNE parameters were set to 30 for perplexity, 12 for early exaggeration, and 200 for the learning rate. For UMAP the normalized MFI expression table for immune cell markers and cells classified as immune cells within ROIs from each QPTiff image were scaled using Scanpy ^119^ with max_value set to 10. Scanpy was used to compute PCA coordinates which were used for batch correction using Harmony ^122^. A neighborhood graph of observations was computed with scanpy and UMAP was plotted with Scanpy and Rapids.

### Single Cell Spatial Profiling of HGSOC with Akoya Fusion

The spatial interactions between different cell phenotypes were quantified using the Cellular Neighborhood method ^123^. A ‘window’ was captured consisting of k nearest cells for each of the cells in all tissues from different patients as measured by euclidean distance based on the X/Y coordinates. Then, for each cell, the cell-type percentages in its neighborhood were calculated. This step produced a neighborhood matrix where cells were characterized by the percentages of cell types in their local neighborhoods. K-means clustering was then performed on all cells based on these percentages with k = {5, 10, 15, 20, 25, 30, 35, 40}. The optimal number of cells within ‘windows’ and cellular neighborhoods were determined as 5 and 15 based on the highest silhouette score (**Supp Figure 8C**). The average neighborhood percentages were also calculated for each cellular neighborhood and plotted on a heatmap (**Supp Figure 8E**).

### Spatial proximity analysis

Spatial proximity analysis was performed by measuring the pairwise distances between different cell phenotypes. This method effectively maps the positions of cell phenotypes relative to one another by calculating the average distance from the nearest cells of type “B” to a target cell of type “A”. Specifically, for each cell of type “A”, the 10 nearest cells of type “B” were identified, and their average distance to the cell of type “A” was computed and attributed as a key feature of cell “A”. Following this, both the mean and median values of this characteristic are determined for all cells of type “A” and used as a new characteristic associated with the sample. Analysis of Variance (ANOVA) was then performed to compare, based on the mean pairwise distance values, between HRD and HRP samples within each group, i.e. Primary or Recurrent (**Figure 6A-E**).

## Discussion

In this study we profiled the proteome of paired chemo-naive and chemoresistant tumors and used data generated on the epigenome and transcriptome of the same tissues. Differences were identified in the tumor immune landscape of high grade serous ovarian tumors based on homologous recombination deficiency. Spatial proteomic profiling confirmed our findings and provided insight into the potential mechanism by which patients with HRD tumors have increased overall survival post diagnosis. The patient population in our study contained a relatively higher proportion of patients with germline *BRCA1* or 2 mutations than in other published studies. Protein expression changes in high-grade serous ovarian cancer tumors were primarily driven by homologous recombination status, as evidenced by differential expression analysis, consensus clustering and identifying modules of co-expressed proteins. Moreover, there were shared changes in the proteome of chemo-naïve and chemo-resistant tumors for individual patients, indicating that mechanisms for chemoresistance already existed in tumors before they were exposed to chemotherapy. These are rare samples to have access to. However, one of the study’s main limitations was the patient sample size. Separating the samples into different clinical groups based on the recurrence and HRD status further reduced the overall sample numbers per group.

Many studies have utilized label-free proteomic approaches to study chemoresistance and progression of ovarian cancer. The label-free DDA-MS based proteomic method identified a 67-protein cell line signature that discriminated predominantly epithelial and mesenchymal HGSOC tumor clusters^40^, discovered CT45 as a prognostic factor linked to disease-free survival^25^, and found NNMT to be a crucial metabolic regulator of cancer-associated fibroblast differentiation and cancer progression in the stroma of HGSOC, making it a prospective therapeutic target^24^. Recently, TMT-based LC-MS/MS analysis identified a 64-protein signature that predicted chemo-refractoriness in a subset of HGSOCs with high specificity ^51^. Compared to previously published ovarian cancer studies discussed above, the number of proteins quantified per tumor in our cohort was lower, which can be attributed to the difference in the sample acquisition method on the mass spectrometer. Despite the low protein numbers quantified, the data-independent acquisition method used in our study allowed for faster sample preparation of a relatively large number of clinical samples with fewer batch effects, more reliable quantification of proteins, and the possibility of re-interrogation of data over time ^89,124^. Mass spectrometry based proteomics can discover new therapeutic targets, as has been shown by the studies described above. But supplementing proteomic data with genomics or transcriptomic data can bring orthogonal validation from multiple analyte types examined on the same sample and enhance the comprehensiveness and accuracy of studying disease biology.

Previous studies have identified an association between HRD status and immune infiltrates in HGSOC ^80^. BRCA1 mutated tumors showed an immunoreactive molecular subtype with intratumoral T cell infiltration ^14,80^, increased CD8+ TILs ^15^, and increased STING and VEGFA levels ^12,16^. We found that in dense neoplastic regions of HGSOC tumors with germline *BRCA1/2* deleterious variants, there was an increased percentage of inflammatory cells and a higher percentage of necrotic and connective cells compared to tumors with no genetic predisposition to HGSOC. Our spatial analysis showed significantly higher levels of STING positive tumor cells in samples with germline *BRCA1/2* mutations. STING positive tumor cells were significantly spatially closer to endothelial cells in HRD primary tumors as compared to HRP primary tumors. Proteins and genes significantly upregulated in HRD tumors were enriched in immune pathways with similar pathway enrichment in an independent HGSOC proteomic data set ^53^. TCR-seq analysis revealed that homologous recombination deficient tumors had a more shared T cell CDR3 repertoire than homologous proficient tumors, exemplified by another independent HGSOC data set ^5^. Shared clonotypes across primary and recurrent HRD tumors from a few patients mainly drove this shared T cell repertoire. HRD tumors also had a significantly higher number of neoantigens than HRP tumors which can be potentially attributed to a higher tumor mutation burden in HRD tumors. Recent research shows that long-term HGSOC survivors had notably more neoantigens and a higher tumor mutation burden compared to those with shorter survival times, highlighting a strong link between neoantigen count and survival. The study used computationally estimated immune cell densities to identify clusters of primary tumor samples. Two immune clusters with the highest neoantigen counts were enriched in plasma cells, activated CD4 T cells, M1 macrophages, and resting NK cells, whereas the cluster with the lowest neoantigen counts had an abundance of resting mast cells and dendritic cells^125^. Another study found a strong positive correlation between strong-binding neoantigens and mutation load in HGSOC ^126^. Tumor cells exhibiting a higher mutational burden have been associated with increased levels of immune infiltration and activity ^127–129^. Similar observations in pancreatic ^130^ and breast cancer ^131^ studies suggest that homologous recombination deficiency may have an effect on immune activation across various cancer types.

Cancer cells with high levels of chromosomal instability utilize the upregulation of inflammatory pathways for metastasis ^132^. PARPis exacerbated the upregulation of the inflammatory state in BRCA1 deficient ovarian cancer cells ^17^. In our study, chemotherapy caused a significant increase in the number of immune cells in HGSOC tumors and tumor microenvironments, irrespective of their HRD status. These immune cells were clustered with other immune cells based on our nearest neighbor analysis. Particularly, HRD tumors had significantly higher number of stromal and necrotic cells near immune-rich neighborhoods after chemotherapy while there was no significant difference in proximity of immune clusters to other cell types after chemotherapy for HRP tumors. Thus, chemotherapy caused an increase in the number of immune cells in HGSOC tumors and tumor microenvironments, and homologous recombination status drove their localization to different compartments of the tissue.

Treatment options are limited for all HGSOC patients, where most patients receive Platinum and Taxane after primary diagnosis. Recently PARPis have been recommended for HGSOC patients with HRD ^10^. We sought to identify proteins involved in immune-related pathways that may be effective therapeutic targets in combination with immunotherapy. One of the top 5 differentially upregulated proteins in HRP tumors was ERAP2, while ERAP1 was upregulated in recurrent HRP compared to recurrent HRD tumors. ERAP1/2 have non-overlapping functions involved in peptide trimming for presentation to immune cells via MHC-class I molecules. Significant upregulation of ERAP1 in HRP tumors, its negative correlation with neoantigen count, and its role in immune suppression in other cancers make it an attractive therapeutic target alongside immunotherapy for patients with homologous proficient tumors. Potential drivers of chemoresistance need to be identified for chemotherapy to be effective in HGSOC tumors. With the differential expression analysis between chemo-naïve and chemoresistant tumors, we identified potential novel therapeutic targets for chemoresistance in HGSOC. Proteins upregulated in recurrent tumors like CD5L, CILP, and TINAGL1 have been predicted to be secreted in extracellular space and can be further studied as potential biomarkers of chemoresistance.

Spatial analysis revealed significant differences in cellular abundances between chemo-naive and resistant tumors irrespective of HRD status. Cytotoxic T cells were significantly higher in chemo-naive tumors while T helper cells were significantly higher in chemoresistant tumors. The difference in levels of Tregs after chemotherapy was dependent on HRD status where recurrent HRD tumors had higher levels and recurrent HRP tumors had lower levels of Tregs compared to their respective primary tumors. We observed significantly higher levels of M1 macrophages in primary HRD tumors compared to HRP tumors. These M1 macrophages were in close proximity to almost all other immune cells in HRD tumors. We observed a more coordinated immune response in HRD tumors based on proximity analysis compared to HRP tumors. Tumor cells positive for STING expression were significantly higher in a subset of HRD tumors with germline *BRCA1/2* deleterious variants and were significantly closer to endothelial cells in HRD tumors. On the other hand, dendritic cells were significantly higher in HRP tumors and were in close proximity to almost all other immune cells compared to HRD tumors. Tumor cells positive for ERAP1 expression were significantly higher in primary HRP tumors compared to HRD tumors. The significant increase in immune cells post chemotherapy in HRP tumors but not HRD tumors (data not yet shown; p-value = 0.025) might be why significant differences based on HRD status in spatial proximity analysis were only observed in chemo-naive tumors. Understanding the differences in immune compositions and their functions in HRD and HRP tumors before and after chemotherapy will be crucial to identify appropriate immunotherapeutic strategies for HGSOC patients.

Thus, in this study, we have shown that homologous recombination deficiency and chemotherapy have unique roles to play in immune activation within HGSOC tumors. We have utilized proteomic data in combination with genetic and epigenetic data to identify potential drivers of HGSOC biology, chemoresistance, and immune activation, which can be targeted in combination with other therapies to improve treatment options for HGSOC patients. We further explored the single-cell spatial relationships of immune cells and highlighted key differences in the immune landscape of HGSOC driven by homologous recombination status and chemotherapy.

---

**Supp Figure 1:**
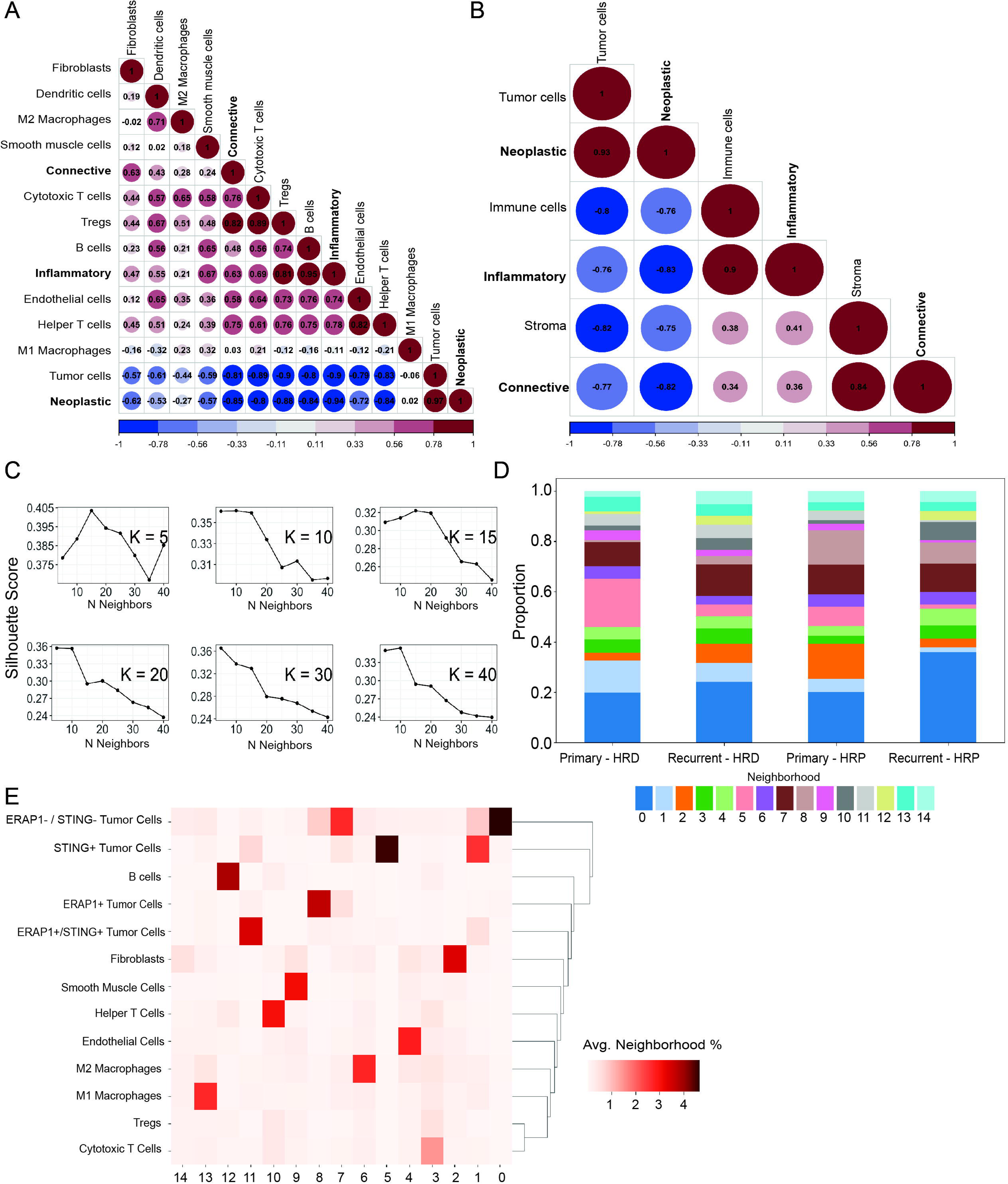
Stromal score based re-classification of samples to mitigate confounding effects of cellular admixtures. A. Hierarchical clustering of Pearson’s correlation coefficients shows clustering of technical replicates and pooled samples with a within-sample replicate mean of 0.95. B. Schematic showing the data sets used to generate the stromal score. B. (Top) Distribution of stromal scores across the re-classified samples. (Bottom) Redistribution of pathology defined samples after re-classification. C. Hierarchical clustering using proteins characteristic to tumor and stromal compartments shows improved clustering of tumor and stromal samples after re-classification. E-F: Volcano plot and Gene set enrichment analysis (GSEA – top 20 pathways) showing proteins significantly upregulated (purple) and downregulated (green) in tumor samples as compared to stromal samples. (threshold: log2 fold change ± 0.5; FDR < 0.01) (Path: Pathology)

**Supp Figure 2:**
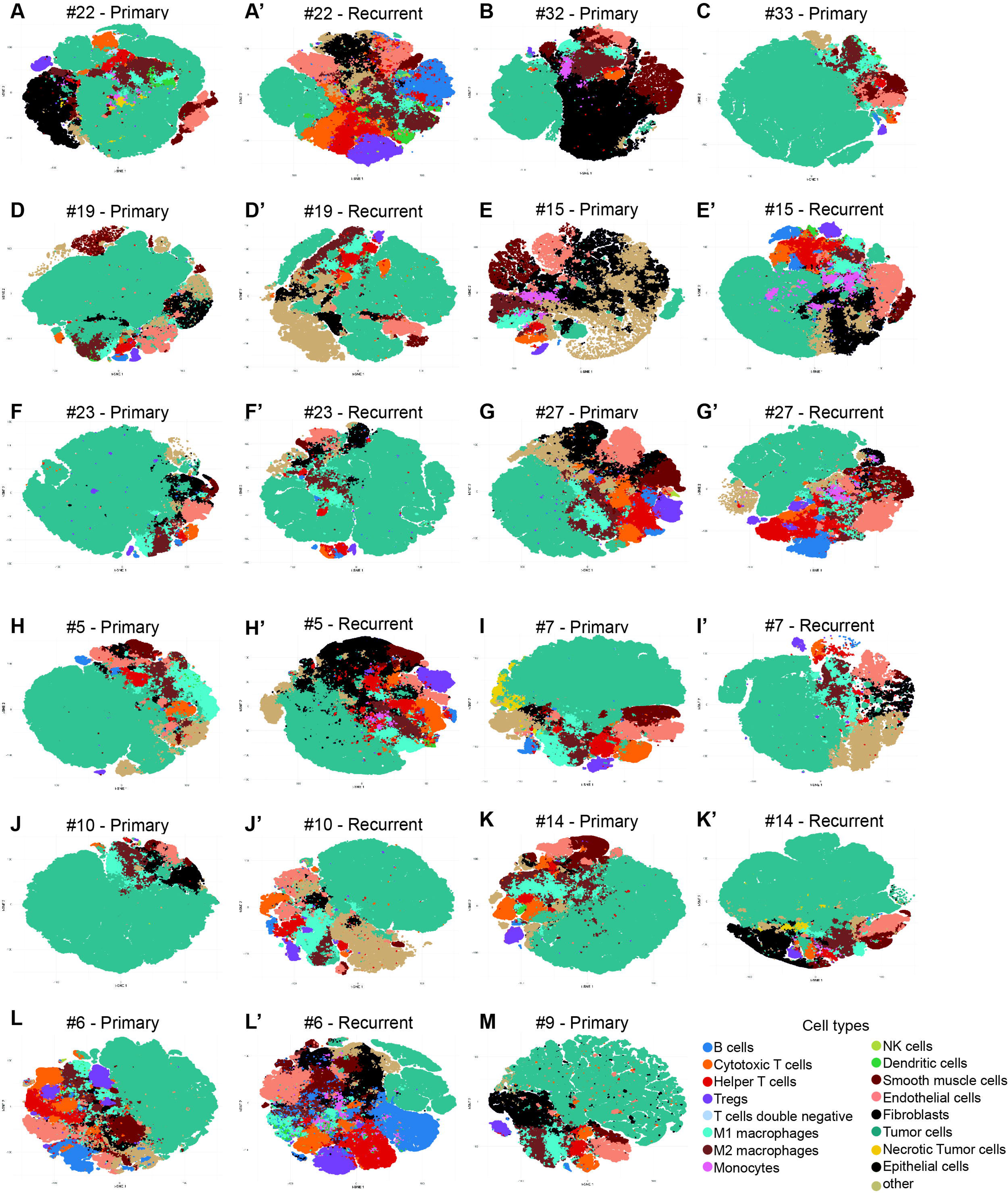
The tumor proteome is stably conserved throughout disease relapse irrespective of the HRD status. A. Protein variability calculated as standard deviation across 59 tumor samples against mean protein intensity shows high variability of frequently altered proteins in ovarian cancer (purple) and low variability for proteins associated with ribosomal biogenesis (green). B. Protein and gene correlation histogram with positively correlated protein-gene pairs shown in purple. A. Top 5 enriched pathways for proteins in decile 1 (most abundant) and 10 (least abundant) with proteins ordered from high to low abundance on the x-axis: GO term enrichment analysis - Biological processes. B. Protein variability calculated as standard deviation across 59 tumor samples against mean protein intensity shows high variability of frequently altered proteins in ovarian cancer (purple) and low variability for proteins associated with ribosomal biogenesis (green). C. Histogram showing correlation between protein and gene expression. Protein expression correlated well to gene expression (measured by total RNA-Seq) with a median Spearman correlation of 0.34 D-E. Protein expression-driven principal component analysis shows heterogeneity among paired primary and recurrent tumors. Six representative patients are shown either clustering (D) or farther apart along the first and second principal components (E). F. Intra-patient Euclidean distance is significantly shorter for paired primary, recurrent samples from the same patient than inter-patient distance, irrespective of HRD status.

**Supp Figure 3:**
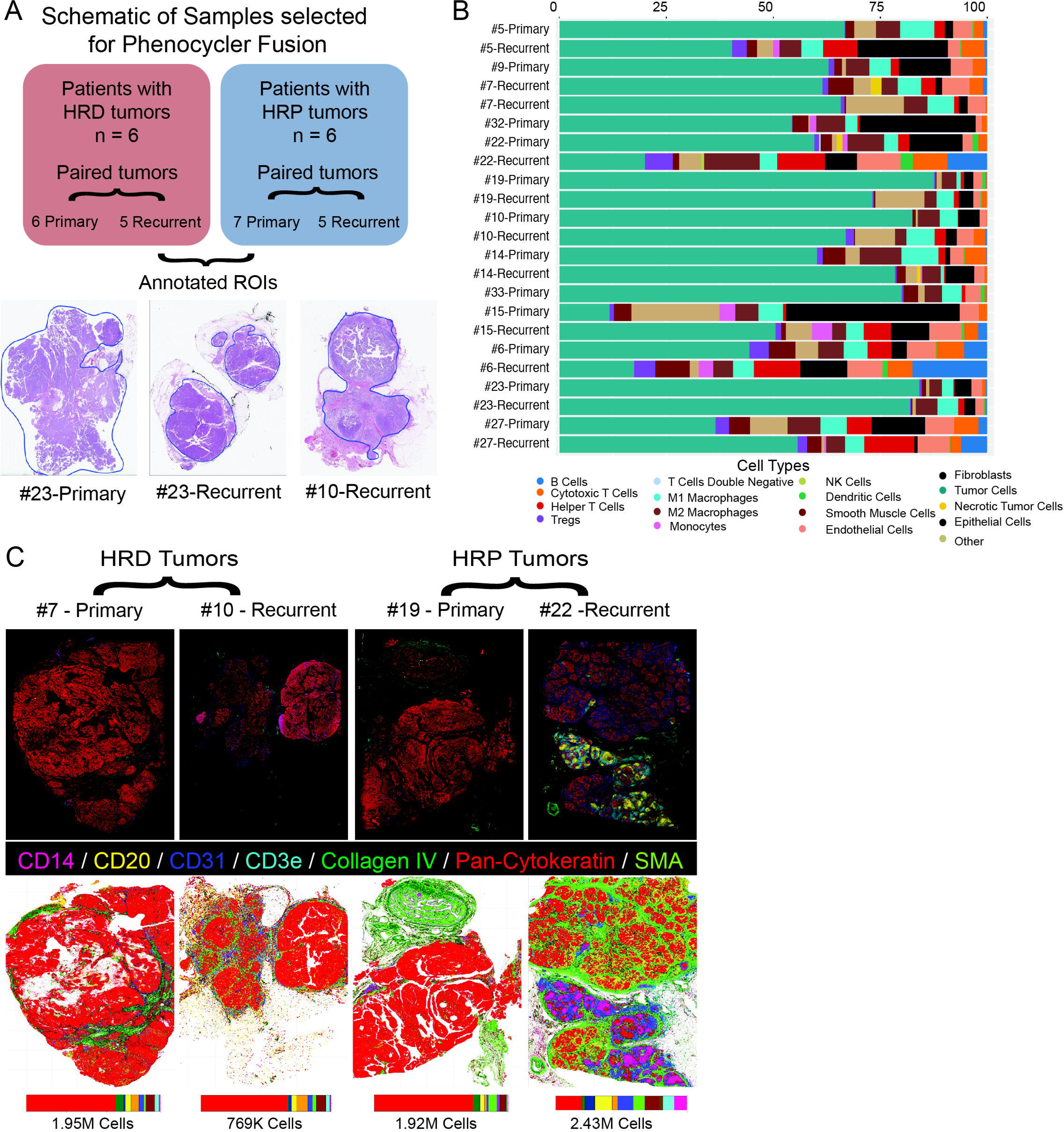
Differential expression analysis between primary and recurrent tumors. A-B. Volcano plot and Gene set enrichment analysis (GSEA) showing proteins significantly upregulated (orange) and downregulated (purple) in primary tumor samples as compared to recurrent samples. (threshold: log2 fold change ± 0.5; FDR < 0.01)

**Supp Figure 4:**
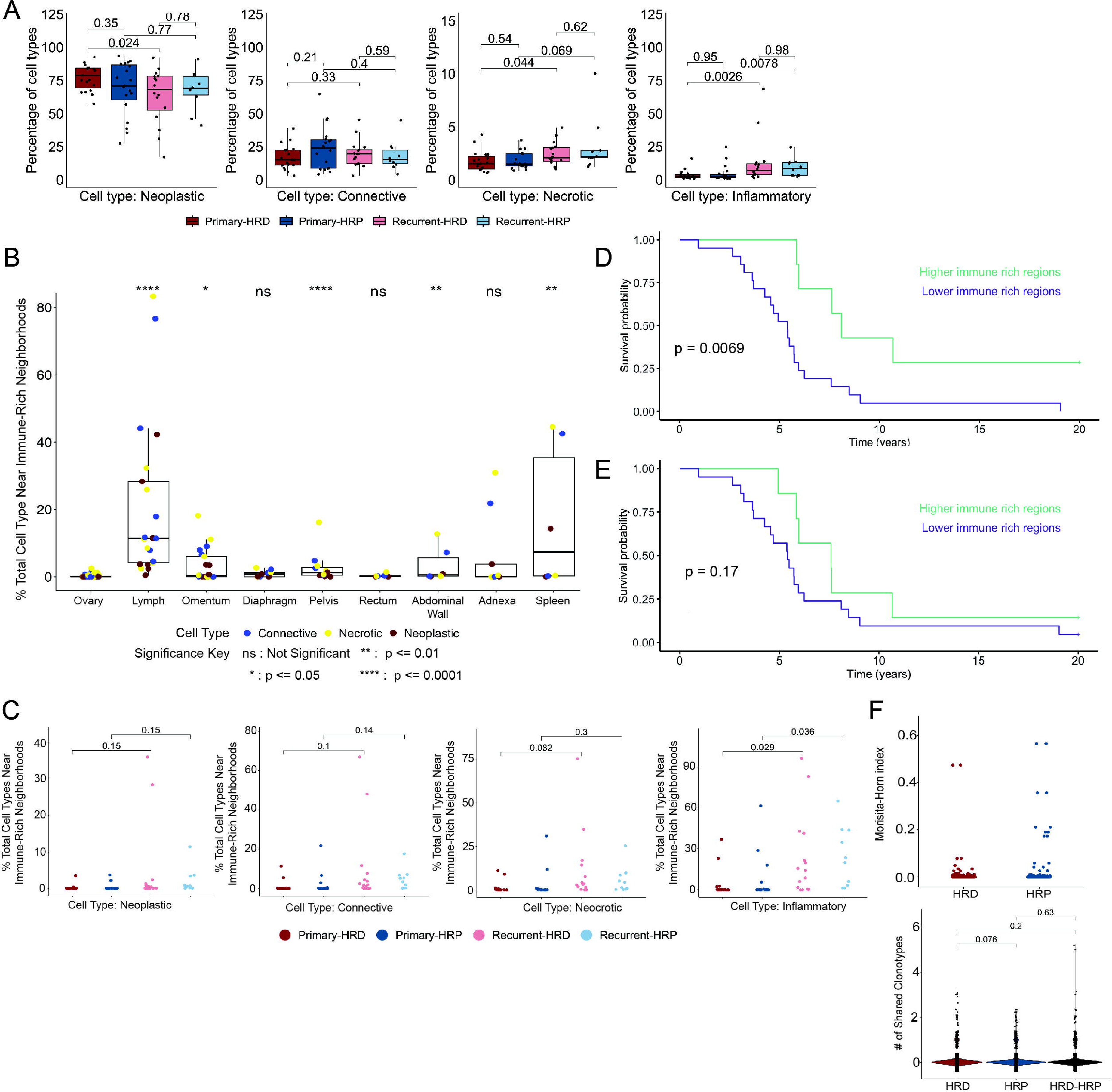
Homologous recombination deficiency drives molecular subtypes identified in HGSOC tumors A. WGCNA modules derived from the CPTAC data correlate significantly with the stromal content of POCROC data compared to other clinical features. (Pearson’s correlation values are shown as numbers in the box, while corresponding p-values shown in brackets) B. Consensus value matrix shows four distinct clusters defined by protein expression profiles of 59 tumor samples from POCROC data. C. WGCNA modules derived from POCROC data correlate significantly with clusters of POCROC data compared to its stromal content. (Pearson’s correlation values are shown as numbers in the box, while corresponding p-values shown in brackets) D. Patients whose primary tumors were in clusters 1 and 4 had a significantly longer survival than those in clusters 2 and 3.

**Supp Figure 5:**
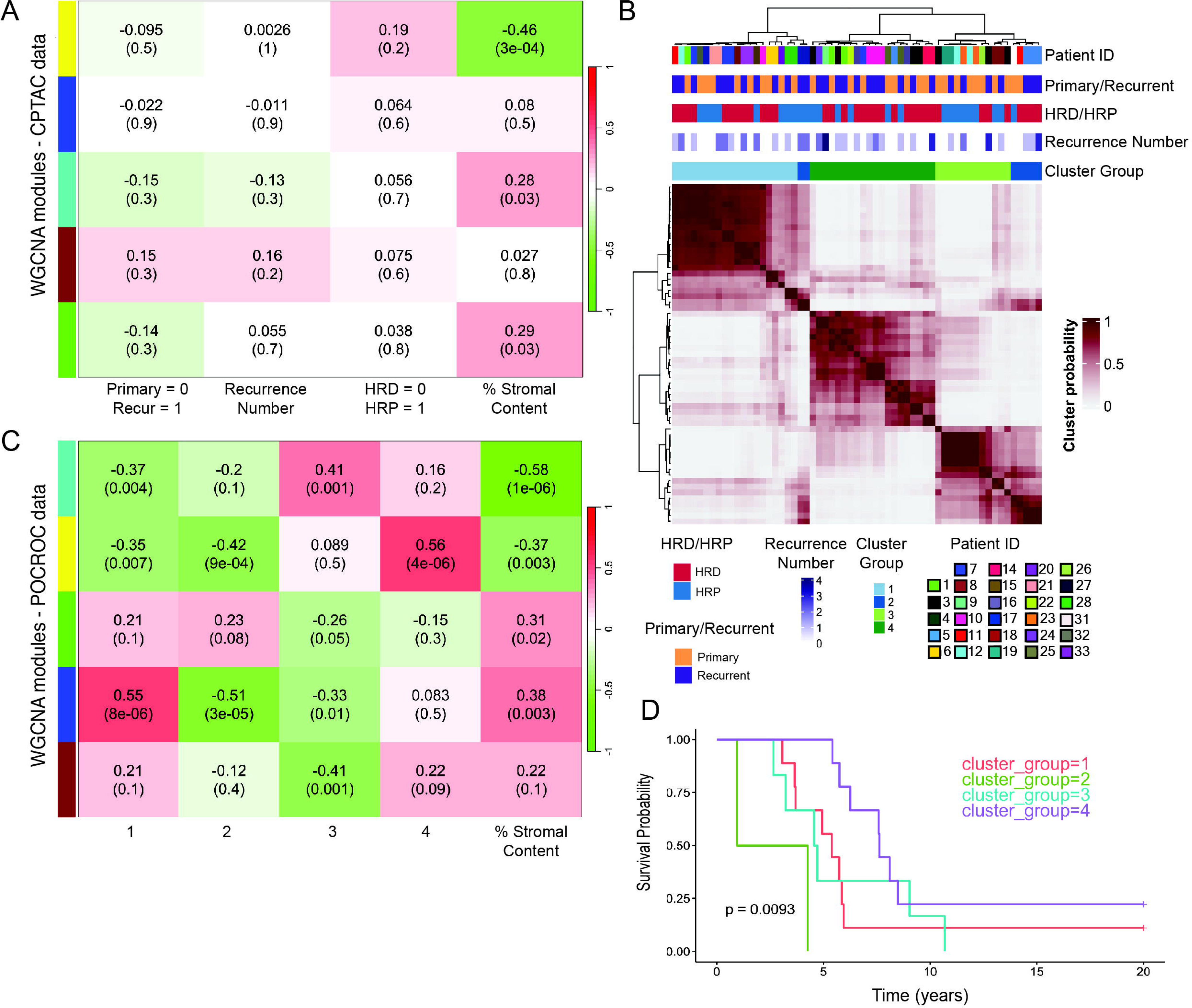
Differences between primary and recurrent proteomes, nearest neighbor analysis, and TCR-seq of HGSOC tumors. A. Boxplot of percentages of cell types in HGSOC tumors and tumor microenvironments grouped by homologous recombination and recurrence status. B. Cells from tumors in the lymph, omentum, and pelvis were significantly closer to immune-rich regions than cells from tumors in the ovary. C. Percentage of cell types near immune rich neighborhoods after ROI selection. D. Increased immune infiltration in recurrent tumors was associated with significantly improved survival in HGSOC patients. E. Survival curve based on immune rich regions proximity to tumor cells after ROI selection. F. Morisita-Horn index calculated from TCR-seq analysis of bulk RNA-seq data from the Bowtell study (Top). The number of shared clonotypes within and across the two groups of samples: HRD and HRP (Bottom).

**Supp Figure 6:**
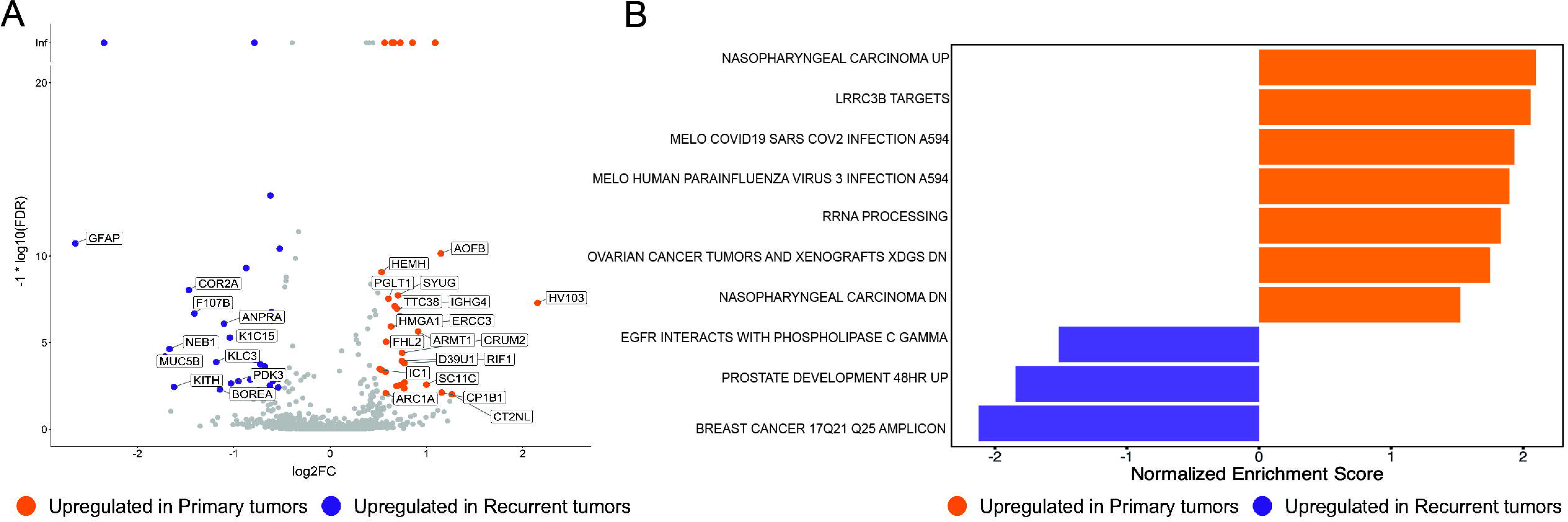
Spatial profiling highlights the pronounced inter and intra-tumor heterogeneity in HGSOC tissues. A. Schematic of samples selected for Phenocycler Fusion with representative manually annotated ROIs shown at the bottom. B. Bar plots showing cellular compositions of the 23 tissues profiled for multiplex staining. C. Single cell spatial maps, accompanied with bar charts, illustrate the differences in the cellular composition and spatial organization of the tumor microenvironment.

**Supp Figure 7:**
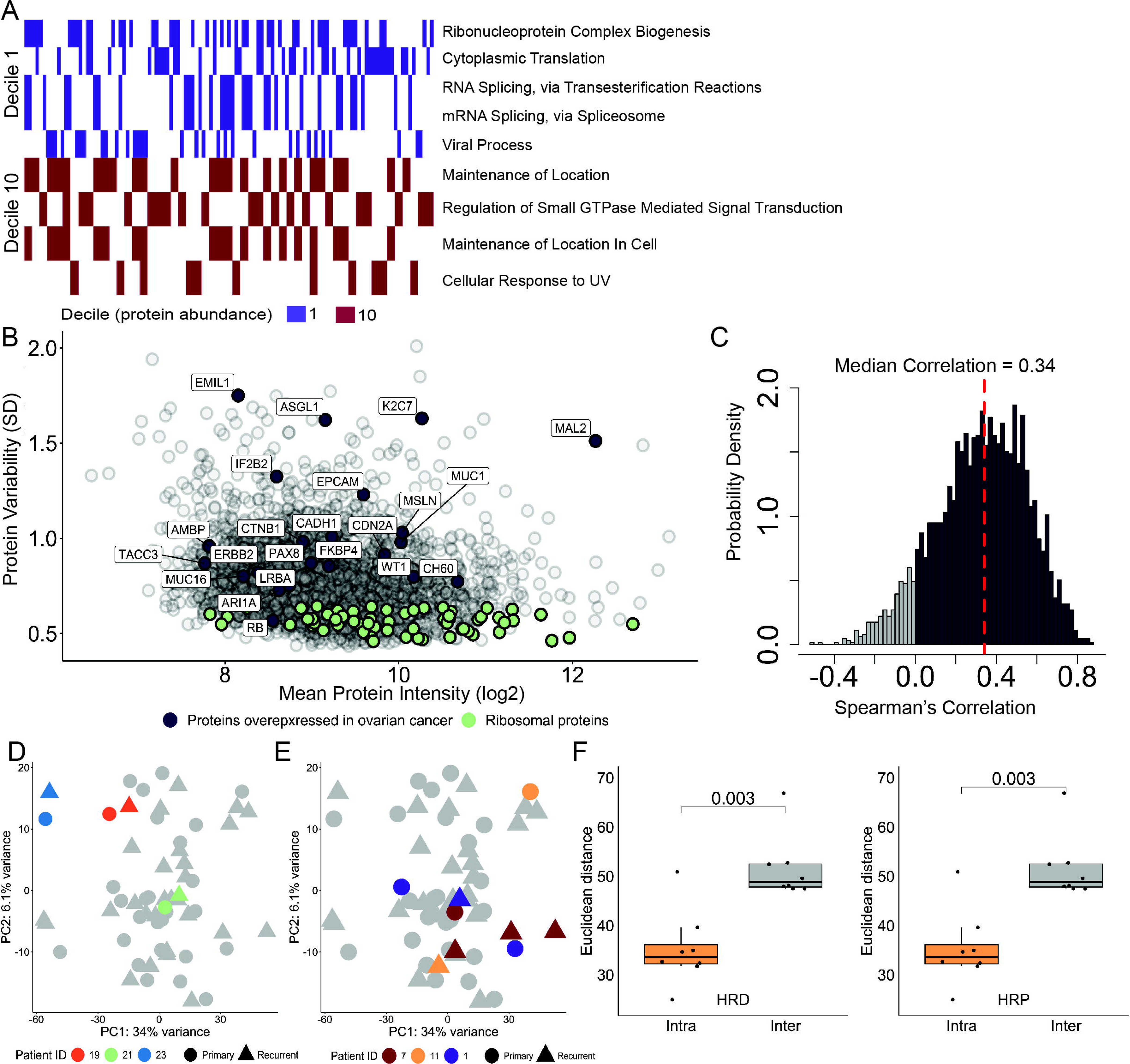
t-SNE representation of all cells of each of the 23 tissues profiled in this study (for paired samples from the same patient, the same alphabet is used but with an added apostrophe)

**Supp Figure 8:**
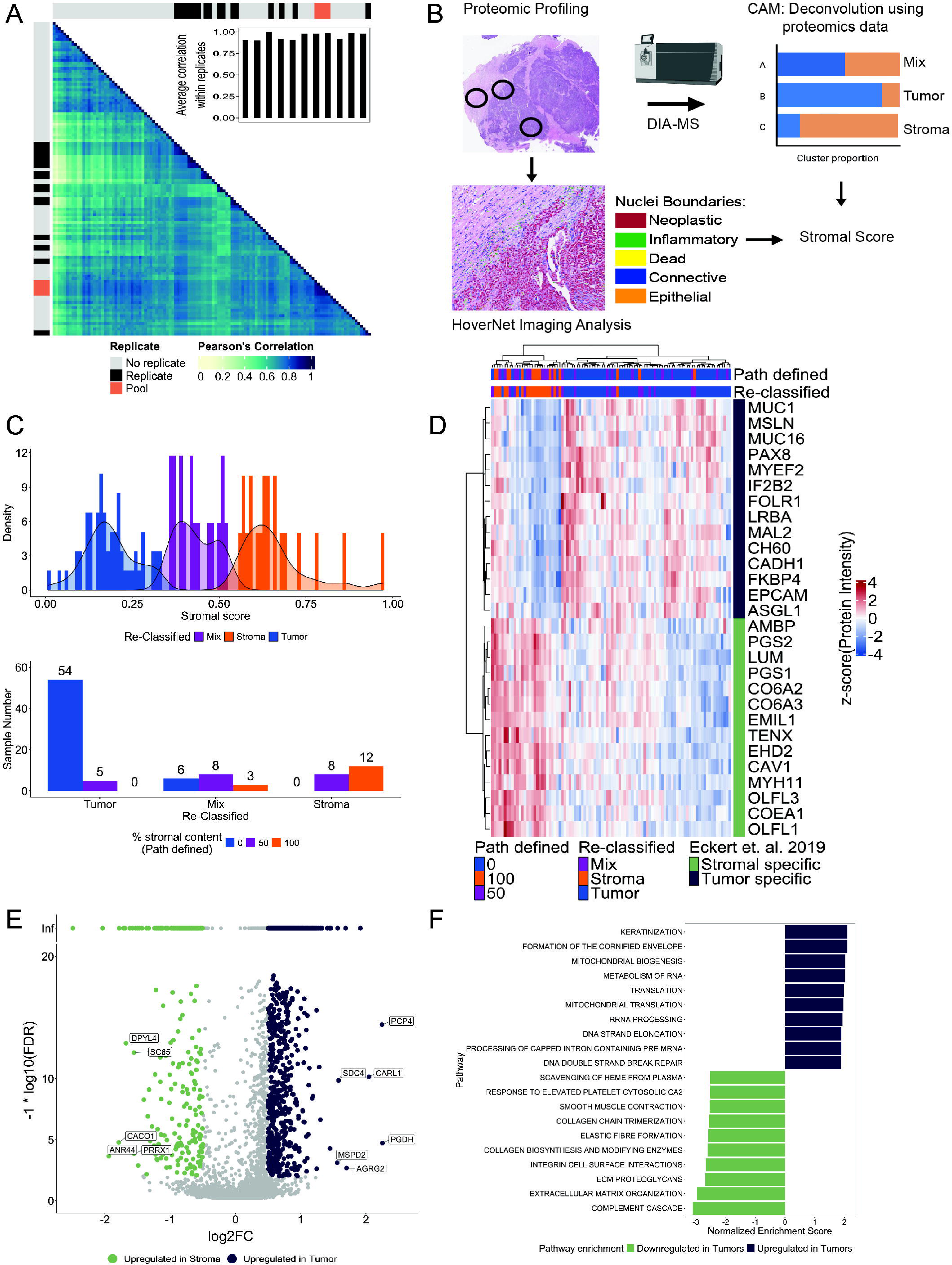
Nearest neighbor analysis shows neighborhood enrichment of cell types based on clinical features of the data. A-B. Hierarchical clustering of Pearson’s correlation coefficients shows clustering of HoverNet annotated cells (in bold) correlated with Phenocycler fusion defined cell types for the 23 samples profiled. C. Silhouette scores for different values of k and n neighbors ranging from 5 to 40 for the nearest neighbor analysis. D. Proportion of neighborhoods across samples based on HRD and recurrence status. E. Heatmap showing relative frequencies of each cell type (y axis) in neighborhoods (x axis) identified from nearest neighbor analysis.

